# Impacts of wind-driven hydrodynamics on the early stages of coral development

**DOI:** 10.1101/2025.05.07.652727

**Authors:** Elizabeth Buccheri, Ananth Wuppukondur, Gerard F Ricardo, Peter J Mumby, Christopher Doropoulos

## Abstract

Coral spawning has evolved to occur during relatively calm hydrodynamic conditions. However, moderate to high surface winds are often present on spawning nights in specific regions, with many unknown impacts to the early life-history stages of corals. Here, we mechanistically examine the sensitivity of fertilisation and larval development to increasing wind speeds for four common species of hermaphroditic corals on the Great Barrier Reef: *Acropora kenti*, *A. millepora*, *A. spathulata*, and *Platygyra daedalea.* Wind intensities relevant to coral spawning seasons were simulated by downscaling location-specific, historical wind speeds to laboratory flumes. Generally, fertilisation success declined as duration of exposure increased and was often lowest at the highest wind intensity equating to 15 knots *in situ*. Embryo damage, deformity, and fragmentation increased with exposure time, but varied across species. Four days following spawning, damaged and fragmented embryos had the highest mortality rates (76-82% and 70-73%) compared to intact embryos (26% and 52%) for *A. kenti* and *A. spathulata,* respectively. Experimental parameters were used to fit a 3D Fluid Dynamics model with outputs confirming that water surface elevations, velocity, and turbulence energy increased by up to 10%, 40%, and 50% respectively as winds intensified, likely explaining the physical drivers of the deleterious effects observed. Overall, results from this study indicate that suboptimal spawning conditions reduce propagule production, which may diminish recovery potential, thus require consideration when planning management strategies to safeguard coral reproduction.

## 1. Introduction

Synchronous mass spawning is the primary mode of reproduction for corals in the Indo-Pacific (Harrison et al. 1984, Willis et al. 1985, Babcock et al. 1986, Harrison and Wallace 1990). The efficiency of synchronisation as a mechanism that maximises fertilisation success is dependent on the interactions among multiple parameters including localised spatial configuration and reproductive timing of fecund adults (Teo and Todd 2018, Mumby et al. 2024, Ricardo et al. 2024), micrometre scale gamete properties of individuals (Thomas 1994), and hydrodynamic conditions that influence gamete interactions (Mead and Denny 1995).

The interaction between hydrodynamic properties like velocity and regime govern gamete condition, mixing (Thomas 1994, Levitan and Young 1995) and ultimate fertilisation success in many spawning invertebrates (Pennington 1985, Denny and Shibata 1989, Levitan, Sewell and Chia 1992, Thomas 1994, Mead and Denny 1995). Pennington (1985) found that fertilisation success was highest for the echinoid *Strongylocentrotus droebachiensis* under moderate hydrodynamic conditions of < 0.2 m s^-1^, and sperm dilution and lower fertilisation occurred when water velocity intensified. Denny, Dairiki and Distefano (1992) observed similar trends for *Strongylocentrotus purpuratus*, where mild turbulence promoted up to 100% fertilisation success in shallow and contained environments. When exposed to higher turbulence, fertilisation declined to as little as 3.9%, despite high gamete concentrations (Denny, Dairiki and Distefano 1992).

Water movement can promote ideal contact times between sperm and eggs up to a critical threshold (Riffell and Zimmer 2007, Crimaldi and Zimmer 2014). Once hydrodynamic forcing gets too high, gamete encounter rates decline (Denny and Shibata 1989, Mead and Denny 1995), owing to the separation of sperm and eggs based on their respective densities (Crimaldi 2012). Invertebrate sperm swimming speeds are often slower than 0.1 mm s^-1^ (Denny and Shibata 1989, Morita et al. 2006), rendering them unable to outswim most water velocities (Denny 1988). Thus, sperm viability and mobility may also be negatively affected by water movement, which further reduces the likelihood of fertilisation and successful larval development (Mead and Denny 1995).

The hydrodynamic environment that hermaphroditic coral gametes are exposed to on spawning nights is different to other spawning invertebrates in the literature. Many invertebrates spawn neutrally or negatively buoyant gametes independently, thus are exposed to more intense mixing due to interactions with the substratum (Denny and Shibata 1989, Mead 1996). Most hermaphroditic coral species have evolved to spawn during neap tides, shortly after the full moon in early summer to take advantage of moderate amplitude tidal conditions (Babcock et al. 1986) that promote gamete retention (van Woesik 2010, Kaniewska et al. 2015). Further, most corals package their sperm and eggs into bundles, which upon release, float to the surface to break and mix with conspecifics (Harrison and Wallace 1990, Mundy and Green 1999). Bundling neutrally buoyant sperm with positively buoyant eggs (Padilla-Gamiño et al. 2011, Kono, Nakamura and Omori 2020) reduces the likelihood of hydrodynamic interference inhibiting fertilisation success by concentrating the sperm and eggs to a two-dimensional space on the water surface and in the upper water column (Miller and Mundy 2005).

Coral egg-sperm interactions usually occur in the top meter of the water column, thus, are more vulnerable to wind generated surface waves and associated hydrodynamics. Despite the notion that corals spawn during very calm climatic periods only, wind speeds have historically exceeded 10 knots during 52% of coral spawning events on the central Great Barrier Reef (GBR) (Heyward and Negri 2012). The intensity of winds, as well as the characteristics of eggs and sperm, will influence the time taken for egg-sperm bundle breakage (Wolstenholme 2004), the degree of gamete movement, and the likelihood of interaction and fertilisation (Mumby et al. 2024). For example, floating eggs at the surface may be transported in different directions and aggregate into clusters based on inertial properties (Denissenko, Falkovich and Lukaschuk 2006, Lukaschuk, Denissenko and Falkovich 2006). The time taken to form aggregations necessary for fertilisation to occur is a function of wave amplitude (Denissenko, Falkovich and Lukaschuk 2006) and Langmuir circulation patterns (Kingsford, Wolanski and Choat 1991), which are both affected by wind speed on a given spawning night.

Wind energy causes shear stress at the surface of the water (Amorocho and Devries 1980), which may cause physical damage to eggs and inhibit fertilisation success and development. The likelihood of egg damage in response to hydrodynamic conditions like turbulence and shear stress has been examined in echinoderms and some fish species (Morgan II et al. 1976, Mead and Denny 1995, Mead 1996, Thomas et al. 1999). Elevated abnormal development and high mortality rates have been observed following fertilisation for *S. purpuratus* in turbulent environments (Mead and Denny 1995). It is believed that echinoderms have evolved extracellular layers, also known as “jelly coats”, in response to the vulnerability of eggs to shear stress (Thomas et al. 1999). Such evolution has not occurred in coral eggs, which remain uncoated and do not contain any protective epidermal layer. Species-specific biological properties influence gamete production (Álvarez-Noriega et al. 2016, Baird et al. 2021), thus gamete tolerance to hydrodynamic stress may vary across taxa.

No studies to date have quantified the influence of surface winds on the highly sensitive stages of fertilisation, embryogenesis, and larval development for spawning corals. Heyward and Negri (2012) examined the influence of turbulent stress on embryogenesis, documenting that fragmented embryos could develop into larvae competent of settlement. However, mechanistic analyses of the effect of wind-driven hydrodynamics on fertilisation and the fate of embryonic development to competency is required to understand widespread effects to demographic processes. Here, we investigated the influence of experimental wind exposure on four species of corals: *Acropora* cf. *kenti* (formerly *A.* “Maggie” *tenuis*), *A*. cf. *millepora*, *A*. cf. *spathulata*, and *Platygyra* cf. *daedalea*. Studies were conducted in enclosed laboratory flume chambers, thus, aimed to replicate windward and shallow reef crest locations, where spawning research is particularly limited due to sub-optimal field conditions. We also applied a Computational Fluid Dynamics (CFD) model to numerically assess hydrodynamic mixing across wind scenarios to help relate our findings to *in situ* environments. Understanding the wind conditions that may optimise fertilisation and the thresholds that result in detrimental outcomes for early development have species- and site-specific implications for management interventions which aim to safeguard coral reproduction and recovery.

## 2. Materials and methods

### 2.1. Quantifying historical wind data

The Coral Spawning Database (Baird et al. 2021) was used to extract the dates and times of spawning from the locations where corals were collected – Orpheus Island (the closest available point of reference for Magnetic Island) and Heron Island – for the most recent decade where data were available from ∼2009 to 2019. The corresponding historical wind data for each spawning record were obtained using the R package ‘dataaimsr’ (Barneche et al. 2021) to compile a range of scalar averaged wind speeds experienced during this season over time. The weather stations where data were collected were Cleveland Bay Weather Station (19.1400 S, 146.9000 E) located off of the eastern side of Magnetic Island (AIMS 2020a), and the Heron Island Relay Pole 7 (23.4600° S, 151.9300° E) located off the southern side of Heron Island (AIMS 2020b).

Four wind speeds of 7, 9, 11, and 15 knots (12.96, 16.67, 20.37, 27.78 km hr^-1^ respectively) were chosen to replicate historical wind patterns most frequently experienced on spawning nights in each study region. Wind data were downscaled to the dimensions of experimental flume chambers based on the principles of similitude (Hughes 1993) by preserving the necessary non-dimensional parameters between the two systems (Hughes 1993). In the case of wind energy, these parameters are the Reynolds number, which represents inertial fluid motion and fluctuations, given by 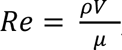, where ρ is the fluid density, *V* is the characteristic velocity, *h* is the characteristic length, and μ is the fluid dynamic viscosity (Denny and Shibata 1989, Rott 1990) and the Weber number (Wb), which represents the forces experienced at the boundary between the surface and the air 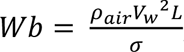, where *V_w_* is the wind velocity, *L* is the length over which fertilisation might occur (or the distance travelled by eggs during fertilisation) and σ is the surface tension of water (Peakall and Warburton 1996). To scale the wind velocities and ensure that the experiments were representative of the environment, we equated the Weber number in the field (referred to as prototype) to that in the laboratory model, (W_b_)_prototype_ = (W_b_)_model_.

### 2.2. Coral collections

Experimental trials were conducted during the 2022 spawning season on the Great Barrier Reef, at the National Sea Simulator (SeaSim) facility in Townsville, Australia in November, and at Heron Island Research Station (HIRS) off the coast of Gladstone, Australia in December. Individual colonies of gravid *Acropora* cf. *kenti*, *A.* cf. *millepora* and *Platygyra* cf. *daedalea* were collected from the reefs surrounding Magnetic Island (19.1385° S, 146.8339° E) in the central GBR in November and individual colonies of gravid *A.* cf. *spathulata* were collected from the reefs on the southern side of Heron Island (23.4423° S, 151.9148° E) in the southern GBR in December.

All corals were identified based on current knowledge of morphological characteristics from the published literature (Wallace 1999, Veron 2000); however, we introduce species with the cf. terminology to acknowledge the uncertainty of species names due to the current and future changes to coral taxonomy (Huang et al. 2011, Bridge et al. 2023). We also acknowledge the possibility of cryptic species creating unknown reproductive barriers for certain coral taxa (Ricardo et al. 2024, Riginos et al. 2024), but it was not logistically possible to conduct genetic analyses to detect cryptic species for this work. Our results show that conspecifics were compatible, thus provide valuable insight into the reproductive mechanisms of these species.

All corals were collected and stored in laboratory aquaria under ambient temperature and light conditions for the spawning period. Individuals were moved into still water isolation containers on expected spawning nights to monitor for setting and spawning (Babcock et al. 1986). At the SeaSim, *P. daedalea* spawned at 18:36–18:42 on the 12^th^ of November, *A. kenti* spawned at 18:10–18:30 on the 13^th^ of November, and *A. millepora* spawned at 20:15–21:05 on the 15^th^ of November. At HIRS, *A. spathulata* spawned at 21:57–22:35 on the 15^th^ of December.

### 2.3. Experimental methods

Laboratory experiments were conducted for each species on independent spawning nights using four recirculating flumes with mixed in-line flow fans to force wind on the water surface. The internal dimensions of the flume chambers were 122.0 x 39.5 x 29.0 cm (length x width x height), with an 80.0 cm long centre dividing wall and semicircular inserts at either end (Fig. 2). Two fans were used for each flume, one in each compartment in opposing corners, with ∼1 m of ducting leading into an explicit region of the tank to direct constant and uniform wind over the water surface and induce a circulating current. Each flume was filled with filtered seawater (FSW) up to a depth of 12 cm to ensure there was at least a 10 cm air column inside the flume available for wind circulation. The 12 cm water depth represented shallow or low tide reef habitats that commonly present physical boundaries to gametes during spawning events, particularly on windward reefs. Tank depth showed no signs of inhibiting processes like bundle breakage which often require surface tension to occur *in situ*. Egg-sperm bundles required longer to dissociate in lower speed wind environments but were still broken by ∼30 mins following initial mixing in most cases.

**Fig. 1.**
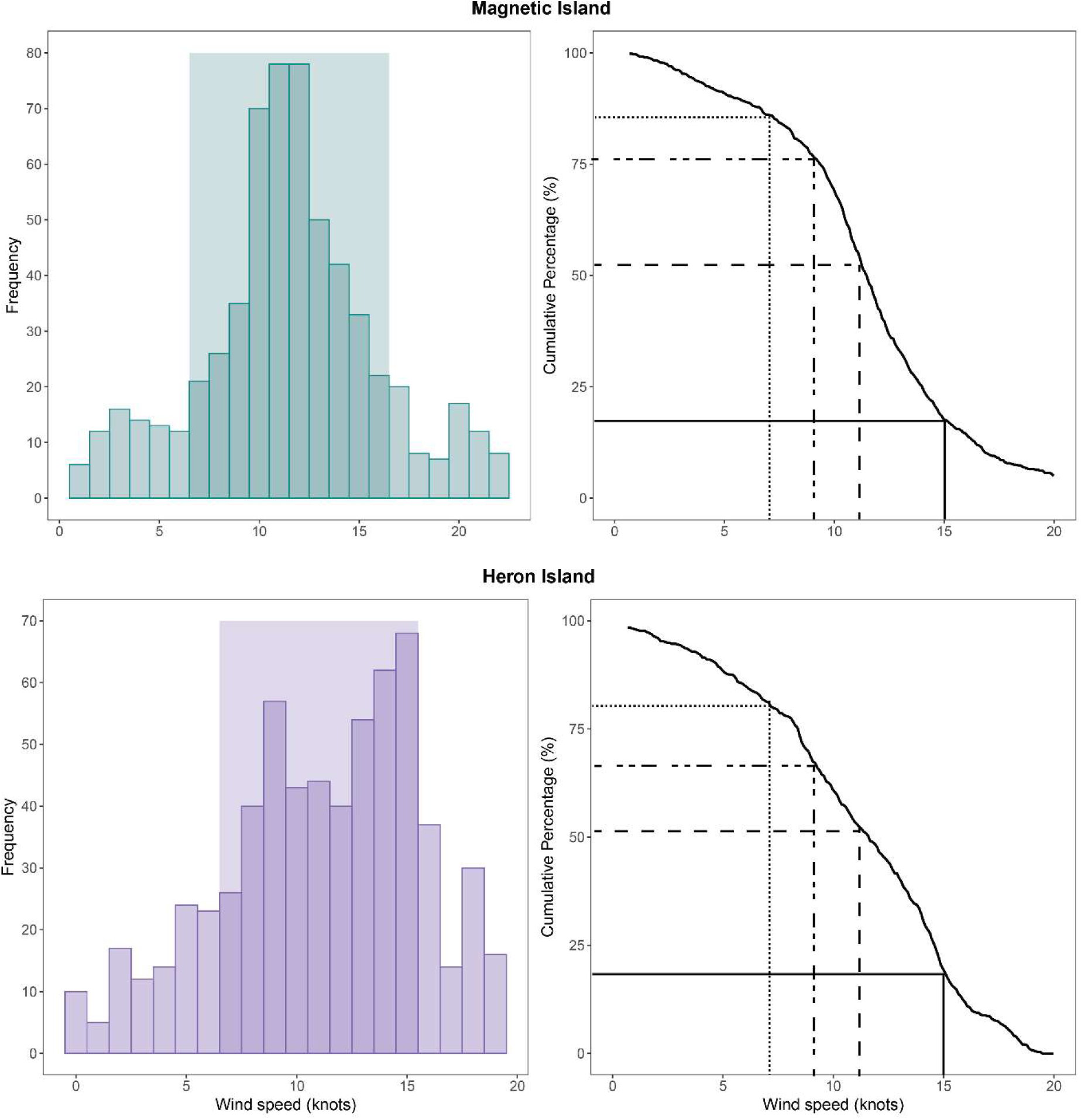
Histograms (left) and exceedance plots (right) depicting the historical wind speeds observed on spawning nights at our two study locations, Magnetic Island in the inshore central GBR (top), and Heron Island in the offshore southern GBR (bottom). Shaded region in the histograms depicts the target wind speed range for experimentation 7 to 15 knots. Dotted, dash-dotted, dashed, and solid lines on the exceedance plots each represent 7, 9, 11, and 15 knot wind thresholds respectively.

**Fig. 2.**
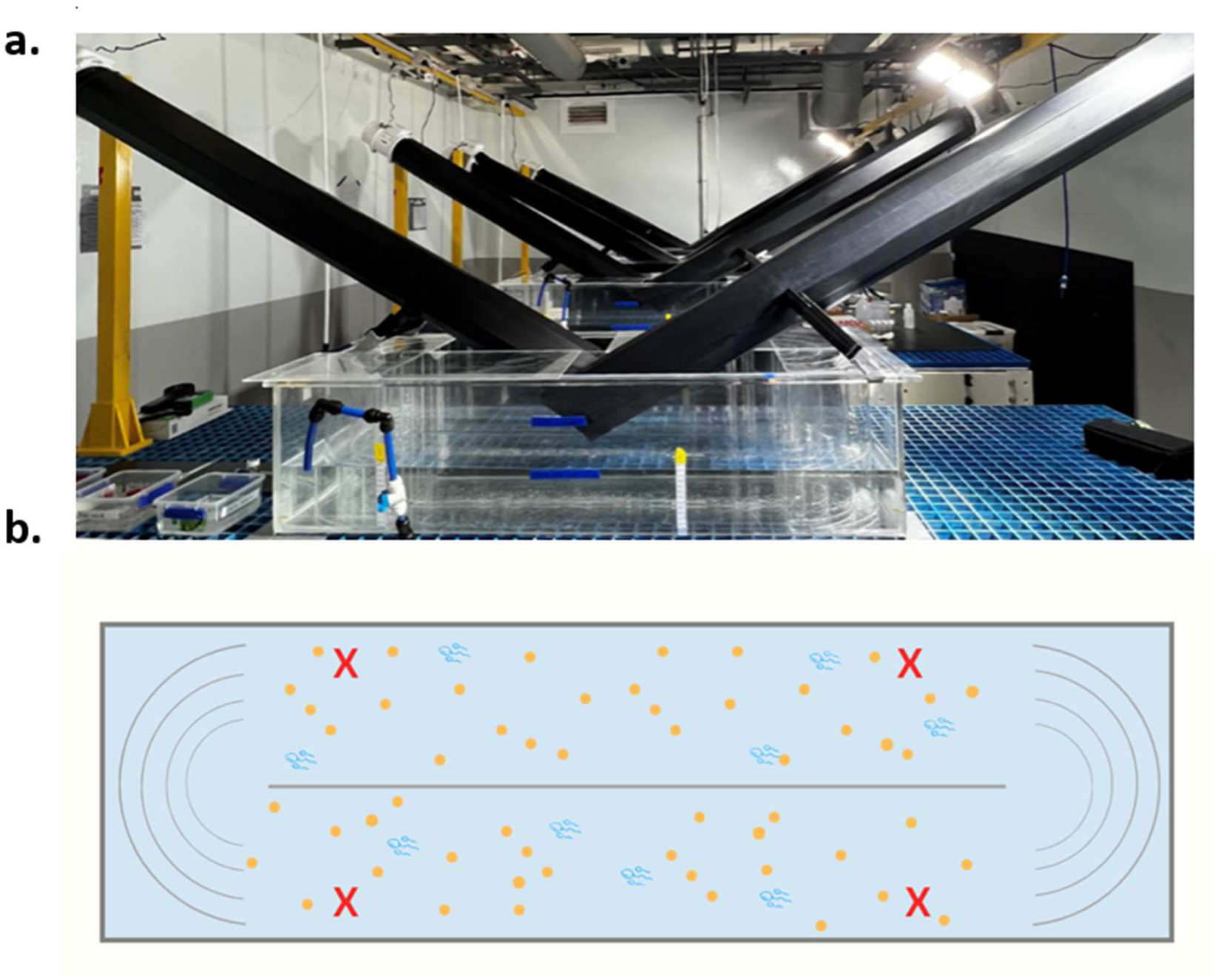
Experimental layout from a profile (a) and top (b) view. Picture of the recirculating flumes with their corresponding fans and ducting systems (a) and a visualisation of the top view of the flume with the divider (providing two compartments) and semi-circular inserts at either end (b). Samples were taken from each corner of the tank every 30 minutes for 3 hours of experimentation.

Prior to experimentation, ∼50 – 125 intact gamete bundles from four individuals were placed into the four respective corners of each flume tank to achieve ∼10^4^ sperm mL^-1^. This sperm concentration was targeted as it is a common inflection point for density-dependent fertilisation thresholds for multiple coral taxa (Ricardo et al. 2016, dela Cruz and Harrison 2020, Buccheri et al. 2023). The quantity of bundles required varied with species due to variability in gamete production across acroporids (∼3–20 eggs, ∼10^6^ sperm per bundle) and merulinids (∼30–250 eggs, ∼10^6^–10^7^ sperm per bundle) (Babcock et al. 2003, Álvarez-Noriega et al. 2016, Madin et al. 2016, Teo et al. 2016, Furukawa et al. 2020). All fans were started simultaneously, and gametes were left to mix with conspecifics. Flume compartments were covered during experimentation to direct the wind over the water’s surface and prevent contamination. Samples were collected from each corner of each flume every 30 min for 3 h following initial mixing. Anemometers were used to measure wind speeds at each point of gamete collection during experimentation to ensure that constant flow was maintained throughout the trials. Thermometers were also used at each sample collection point to track the degree of inevitable wind-driven water temperature declines over time.

After collection, samples were immediately rinsed with sodium lauryl sulfate (SLS) for 10 s and FSW for 20 s in two separate water baths. SLS is a surfactant that denatures proteins in coral sperm (Allen and Hagstrom 1955), thus ensuring experimental contact times were explicit (Buccheri et al. 2023). Samples from each timepoint were left to develop in FSW for ∼3 h from the commencement of the experiment and then fixed in a 4% buffered formalin solution in filtered sea water with 10 g L^−1^ sodium β-glycerophosphate at a ratio of 1:4 fixative to sample to prevent further embryonic development. Samples were scored using a dissecting microscope in the following days to quantify normal fertilisation, damage, deformity, two types of fragmentation, and unfertilised eggs following different wind intensities and exposure periods.

The six conditional categories were distinguished from one another using a consistent scoring rubric (Fig. 3). Embryos were classified as intact if they underwent normal cell division with no apparent disturbances (Fig. 3A). Embryos were classified as damaged if there were obvious large rips or pulls in the cellular tissue (Fig. 3B) and deformed if they exhibited less serious effects of abnormalities like uneven cleavage (Fig. 3C). Fragments were classified as “with divisions” if they were smaller than the normal embryo range for their species and exhibited signs of breakage, while still having two or three cleaved cells intact (Fig. 3D). Fragments “without divisions” were within the fragment size range but were only an individual segment of a cleaved cell (Fig. 3E). Unfertilised eggs were classified if they showed no signs of cleavage and were also not damaged, deformed, or fragmented (Fig. 3F).

**Fig. 3.**
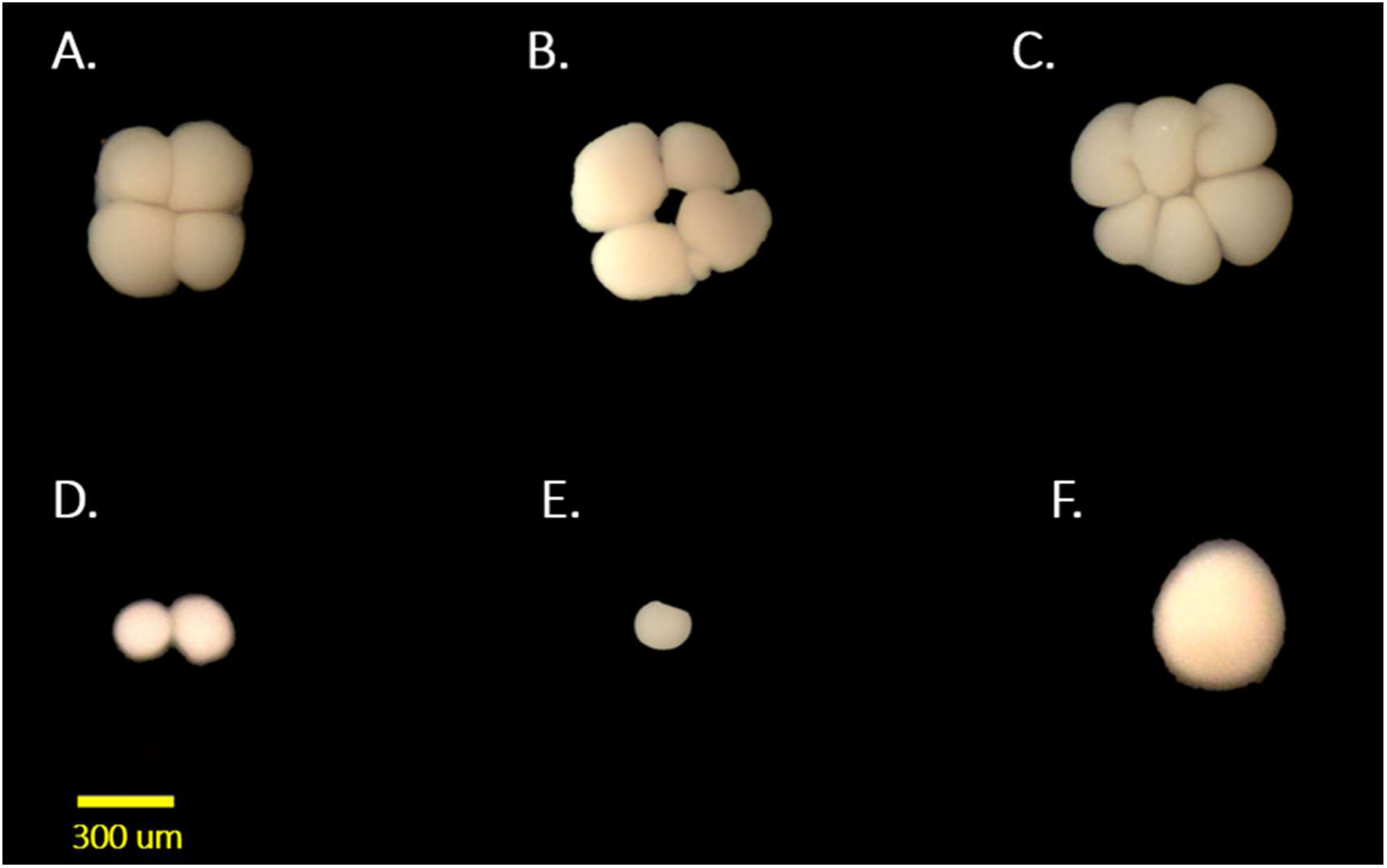
Visual representations of each condition category scored following wind exposure. Intact 4-cell stage (A), damaged 4-cell stage (B), deformed 8-cell stage (C), fragmented 4-cell with intact divisions (D), fragmented 4-cell with no divisions (E), and unfertilised egg (F).

For *A*. *kenti* and *A*. *spathulata*, following the initial 3 h of experimentation on spawning nights, a subset of the remaining embryos from the wind treatments were collected and sorted into 6-well plates with individuals that matched their categorical description, and reared to quantify subsequent survival. Six replicate wells were used for each condition treatment, with 20—50 total embryos per condition depending on feasibility of collection post-experimentation. Embryos were stored in 0.2 µm FSW and water changes were conducted daily to maintain water quality and prevent incidental mortality. Plates were counted at least once a day using a dissecting microscope to determine the number of survivors across condition classes and treatments. Dead embryos were removed, and surviving larvae were tracked until their swimming and searching phase, just before settlement. Rearing was carried out for as long as logistically possible, 4 days following spawning for *A. kenti* and 5 days for *A. spathulata,* which is an ecologically relevant period to capture larval mortality metrics following stress (Okubo et al. 2017).

The time-sensitive nature of spawning experiments and the large scale of the laboratory flumes prevented tank replication of each wind speed treatment for each species. Further, replicating wind speeds for each species across different spawning nights was not possible due to the lack of consecutive spawning events with adequate gametes present to run experiments of this scale. To address this lack of replication, we collected multiple samples from each tank at each time point, which although pseudo-replicates, provided additional data for the analysis. Low replication in large-scale carbonate (Langdon et al. 2003, Albright et al. 2016) or hydrodynamic (Beuzen et al. 2018, Atkinson and Baldock 2020) studies is common when the scale of the experiment precludes multiple chamber or temporal replicates.

### 2.4. Statistical analyses

Alternative Dirichlet regression models were run using the *DirichletReg* package (Maier 2021) in R version 4.3.2 (RCoreTeam 2023), to evaluate the interactive influence of wind intensity and duration of exposure on the six main categories of embryo condition: fertilised, damaged, deformed, fragmented with cell divisions, fragmented without cell divisions, and unfertilised. Dirichlet regression models were used to examine the response variable as the number of embryos across each of the six condition categories due to the covarying nature of the design, rather than evaluating each condition category separately in univariate models that would violate the assumption of independent samples. Model selection and diagnostics were run by examining the fitted and residual outputs from each model and comparing them to the observed data to examine goodness of fit. Probability values were corrected using the Holm method with the *p.adjust* function in the base stats package to accurately conduct hypothesis testing across multiple comparisons in the analysis (Holm 1979).

The *surv* function from the survival package (Therneau 2023) was used to evaluate the influence of the condition of each embryo, and the number of days since exposure to wind on survival over a 4- or 5-day period. A random variable was included with days since mixing nested within the well ID number to account for the repeated measures of documenting each group daily. The two types of fragmented embryos were pooled for the analysis due to lack of adequate replication separately. The *survdiff* and *coxph* functions from the survival package were used to test for statistical differences between the survival rates of damaged, deformed, and fragmented embryos compared to the normal embryos. Plots were created with the *ggsurvplot* function from the survminer package (Kassambara, Kosinski and Biecek 2021) to visualise the results.

### 2.5. Hydrodynamic modelling

Hydrodynamic numerical modelling was implemented using FLOW-3D® HYDRO Version 2023R1 (FLOW-3D 2023) to investigate the degree of change in hydrodynamic conditions across wind treatments and resulting influence on coral eggs and sperm within our experimental flumes (Supplementary materials Table S1).

Computational Fluid Dynamics (CFD) modelling has been used previously to examine wind shear induced water currents, but has focused on unidirectional forcing due to limitations of modelling recirculating environments (Plate 1970, Komori, Nagaosa and Murakami 1993, Zounemat-Kermani and Sabbagh-Yazdi 2010, Druzhinin, Troitskaya and Zilitinkevich 2012). To address this, we applied a novel approach to replicate the wind shear stress on the water surface and the resulting circulation observed in the flumes. “Periodic” boundary conditions were adopted to facilitate movement of fluid and particles leaving the right-hand boundary and being reintroduced into the left-hand boundary, ensuring continuous circulation in the flume, and simulating a recirculating environment.

Four turbulence models were tested, and the most feasible option was selected based on accuracy and feasibility of modelling an experiment of this scale. Further, cylindrical objects were introduced to either end of the modelled flume tank to replicate the eddies and turbulence created by the curved dividers in the physical model. Additional details about model selection, development, and validation are described in the Supplementary materials, Methods.

The primary output parameters from the CFD model included water surface elevation (m) (used as a proxy for wave height), water velocity (m s^-1^), turbulent kinetic energy (m^2^ s^-2^), and turbulence intensity (m^2^ s^-2^). These parameters were used to evaluate the extent of particle mixing and the shear-induced stress experienced by gametes under different wind regimes

## 3. Results

### 3.1. Historical wind data

Historical wind data on spawning nights was similar across study sites, ranging from 0 to 20 knots with an average speed of 11 knots in both locations. However, there were more frequent intense winds on Heron Island compared to Magnetic Island (Cleveland Bay). Wind events at Magnetic Island exceeded 7 knots >80% of the time, 9 knots ∼75% of the time, 11 knots ∼50% of the time, and 15 knots <20% of time on spawning nights. While wind events at Heron Island exceeded 7 knots >75% of the time, 9 knots <75% of the time, 11 knots ∼50% of the time, and 15 knots <20% of the time (Fig. 1).

### 3.2. Fertilisation and embryo development

*Acropora kenti* had the highest and most rapid fertilisation success of all species, with >75% fertilisation at the shortest contact time of 30 min (Fig. 4). Fertilisation for *A. kenti* was consistent in the 7-knot wind treatment but gradually declined after 90 min of exposure in the higher 11 and 15 knot wind treatments. Embryo damage and deformity increased as duration of exposure increased in the 15 knot and 11 knot wind treatments, respectively. No statistically significant trends were observed following p-value correction (Supplementary materials Table S2).

**Fig. 4.**
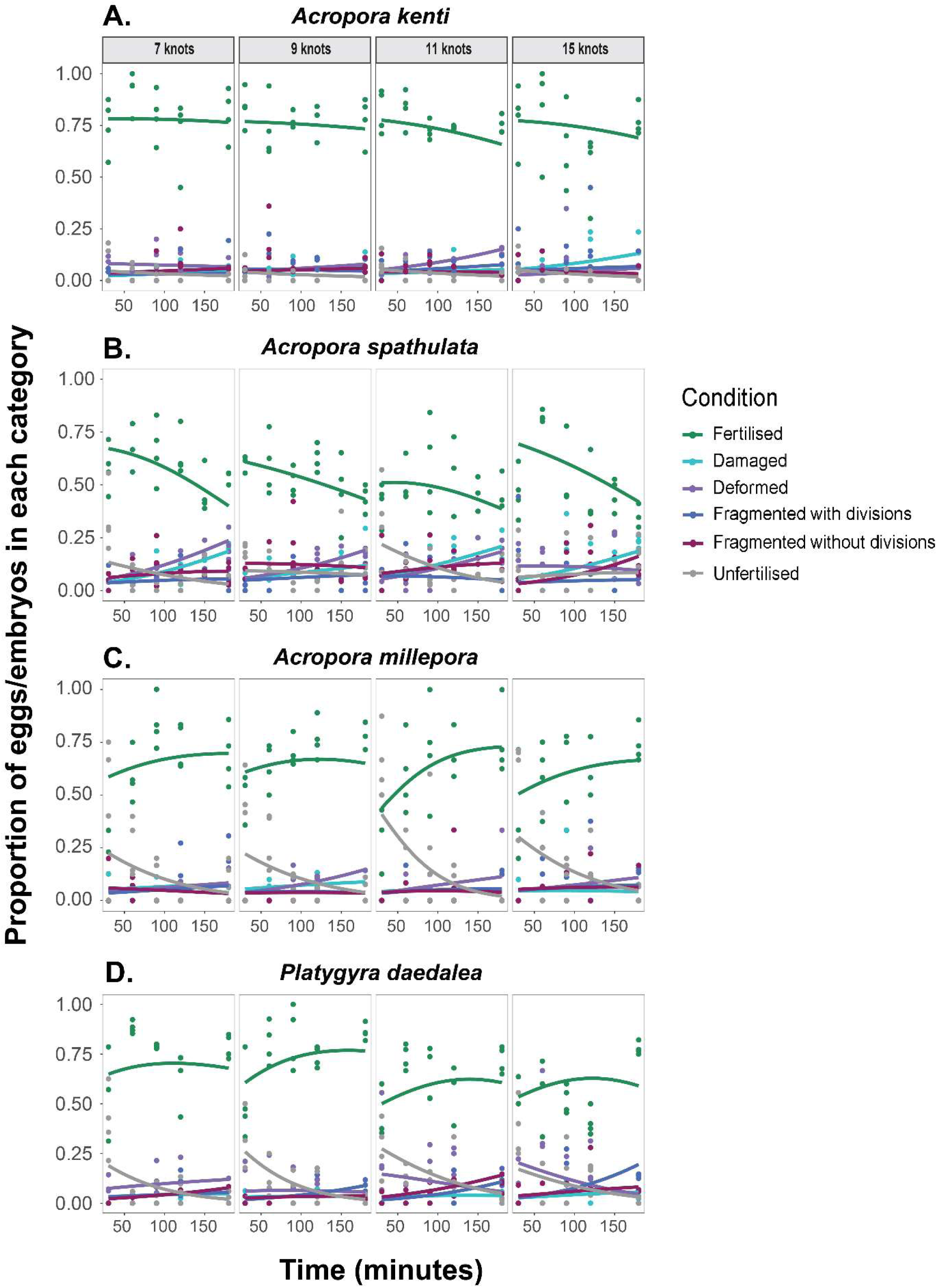
Results from alternative Dirichlet regression analyses evaluating the effect of wind speed and duration of exposure on fertilisation and embryo condition. Each row delineates a species, and each column delineates wind speed treatments. Colours represent condition categories that were scored during the analysis. Each dot represents the raw data from the experimental trials over time for each species, and smoothed lines represent the model prediction for each condition category.

*Acropora spathulata* exhibited similar but more severe trends in fertilisation, with declines of at least 20% from the shortest (30 min) to longest (3 hr) contact times in most cases (Fig. 4). Damage, deformity, and fragmentation were highest for *A. spathulata* compared to all other species as time progressed. After three hours, there was substantial deformity (20-25%) in the 7, 9 and 11 knot wind treatments, and damage (20-25%) in the 7, 11, and 15 knot treatments. Duration of exposure was a statistically significant predictor of both deformity (*p=0.009*) and damage (*p=0.007*) (Supplementary materials Table S2).

Trends for *A. millepora* were less pronounced than *A. kenti* and *A. spathulata,* with fertilisation increasing gradually over time (Fig. 4). Fertilisation success was <50% at the shortest contact time of 30 min and the percentage of unfertilised eggs was 25%. Damage and fragmentation were minimal (5-10%), and deformity was highest (10-15%) in the 9 and 11 knot treatments after 3 h. No statistically significant trends were observed following p-value correction (Supplementary materials Table S2).

*Platygyra daedalea* exhibited similar relationships to *A. millepora*, with slight increases in fertilisation over time. Damage was minimal (<5%), and fragmentation was more common (10-20%), especially in the 11 and 15 knot treatments at the longest contact times (3 h; Fig. 4). Deformity was highest in the first hour of exposure in the 11 and 15 knot treatments and steadily increased over time in the 7-knot treatment. Duration of exposure was a significant predictor of fragmentation without divisions (*p=0.010*) and showed a trend towards statistical significance for fragmentation with divisions (*p=0.074*) and deformity (*p=0.069*) (Supplementary materials Table S2).

### 3.3. Latent larval survival

For *A. kenti,* survival of damaged, deformed, and fragmented embryos was significantly lower than that of normal embryos (*p<0.001*) (Supplementary materials Table S3, S4). Most normal embryos survived to the swimming stage, with only 26% mortality after 4 d (Fig. 5). In contrast, damage, fragmentation, and deformity during embryogenesis all resulted in relatively high mortality rates from 76 to 82%.

**Fig. 5.**
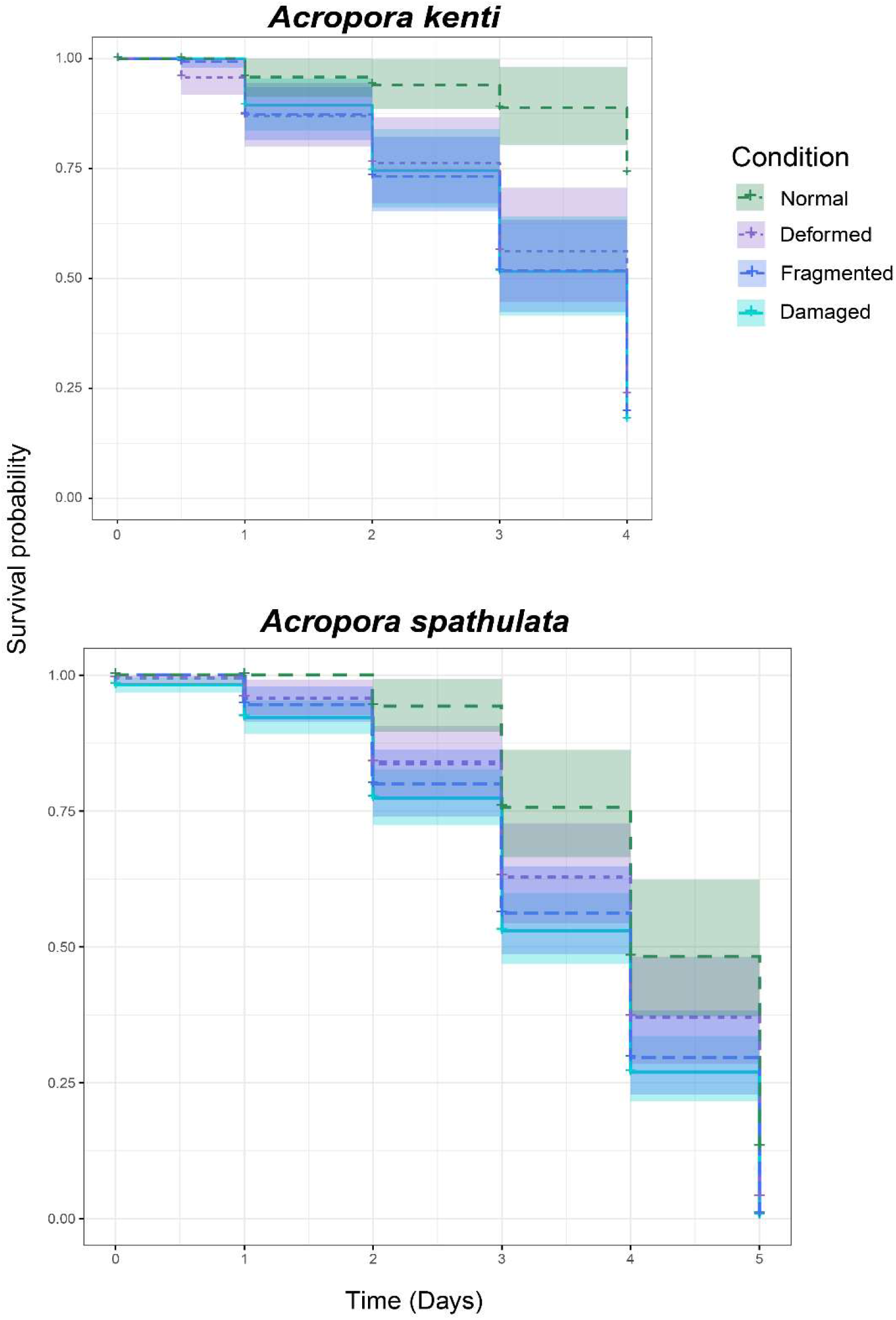
Survival curves depicting the proportion of surviving embryos from each condition category in the days following spawning. Each colour represents a different condition category, and shaded regions represent 95% confidence intervals.

Survival outcomes for *A. spathulata* were comparable across condition groups, with damaged and fragmented embryos having the highest mortality rates and the most significant differences in survival compared to normal embryos (*p<0.001*). Survival rates for deformed embryos were also significantly lower than normal embryos, but trends were less significant (*p=0.010*) (Supplementary materials Table S3, S4). Overall mortality rates were similar to *A. kenti* following 4 days for all impacted embryo conditions (63-73%), but substantially higher after 5 d for normal and impacted embryo conditions (Fig. 5). We observed 51-87% mortality for normal embryos, 63-96% for deformed embryos, and 73-99% for fragmented and damaged embryos after 5 days.

### 3.4. CFD modelling

A total of 15 simulations were performed to evaluate the capabilities of FLOW3D-HYDRO in simulating wind shear induced currents and the resulting dispersion of coral eggs and sperm. Wind speed, turbulence model choice, and cylindrical obstruction implementation all influenced wind-generated hydrodynamic conditions.

The water surface elevations, velocity, and turbulence energy increased by up to 10%, 40%, and 50% respectively as wind speed and time increased (Supplementary materials Fig. S1). Water velocities were highest along the edges of the flumes (Fig. 6). Mean turbulence energy was ∼4.0 x 10^-4^ m^2^ s^-2^ in the lowest wind treatment of 7 knots and reached a maximum of ∼1.6 x 10^-3^ m^2^ s^-2^, compared to the highest wind treatment of 15 knots, that experienced a mean of ∼7.0 x 10^-4^ m^2^ s^-2^ and a maximum of 2.1 x 10^-3^ m^2^ s^-2^. Turbulence intensity increased slightly as wind speed increased, with a mean value of ∼15 m^2^ s^-2^ in the lowest wind treatment (7 knots) and a mean of ∼20 m^2^ s^-2^ in the highest wind treatment (15 knots). Higher turbulent energy and turbulent intensity was experienced at the edges of the compartments, where cylindrical obstructions were placed in the simulations (Fig. 7), and where semi-circular inserts were in the laboratory experiment (Fig. 2). Cylindrical obstructions also successfully promoted lateral mixing, thus ensuring that particles were dispersed across the flume homogenously (Fig. 6). Additional results regarding turbulence model selection and particle dispersion are outlined in the Supplementary materials, Results.

**Fig. 6.**
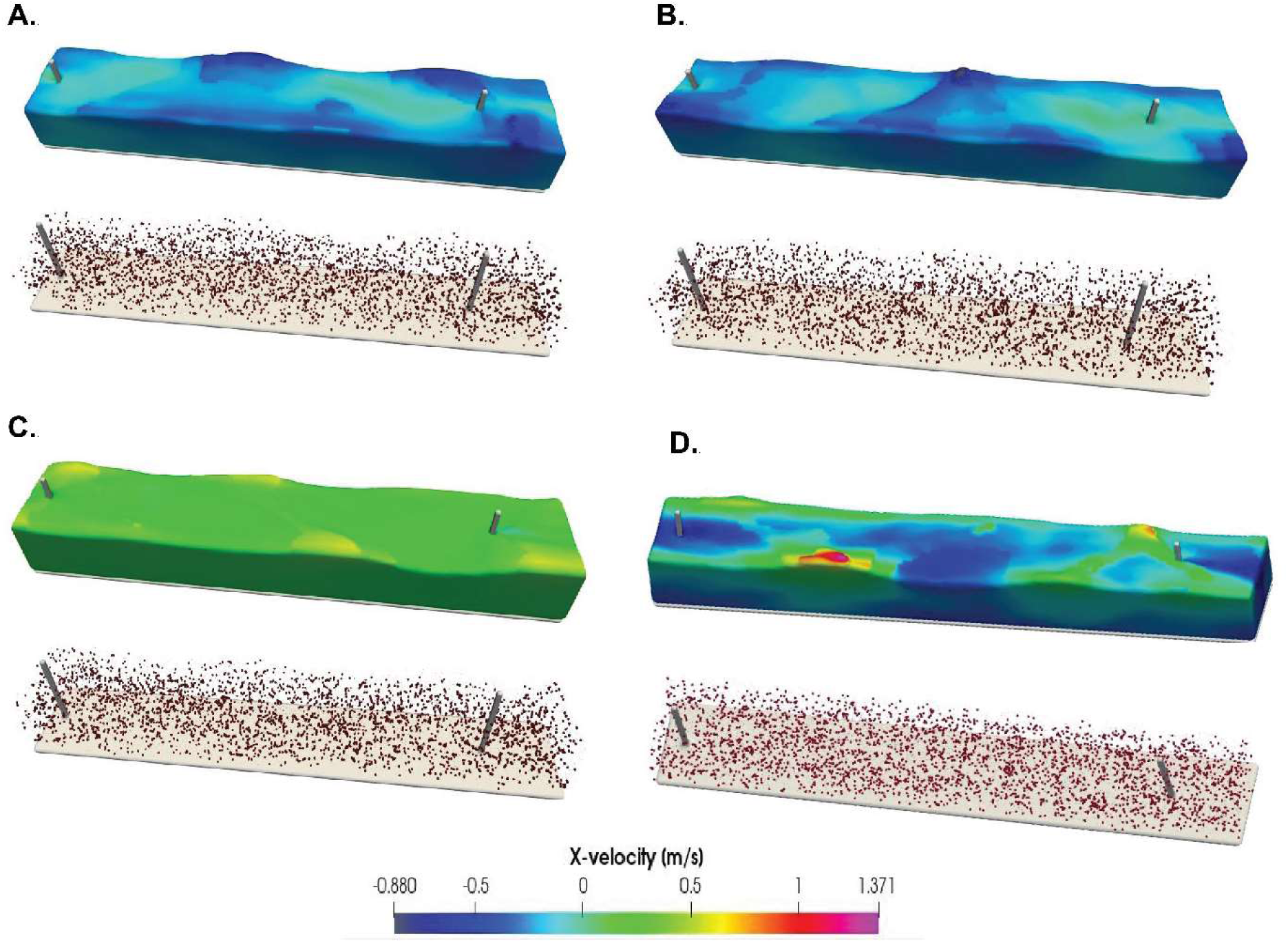
Wind generated current velocities (m s^-1^) in the flume predicted in the simulations for representative examples of (a) 7 knots, (b) 9 knots, (c) 11 knots and (d) 15 knots. Each panel shows the horizontal x-fluid velocity contour map of in the flume (top) and visualises the position of egg (red) and sperm (blue) particles in the fluid after flow stability is achieved in the numerical model (bottom).

**Fig. 7.**
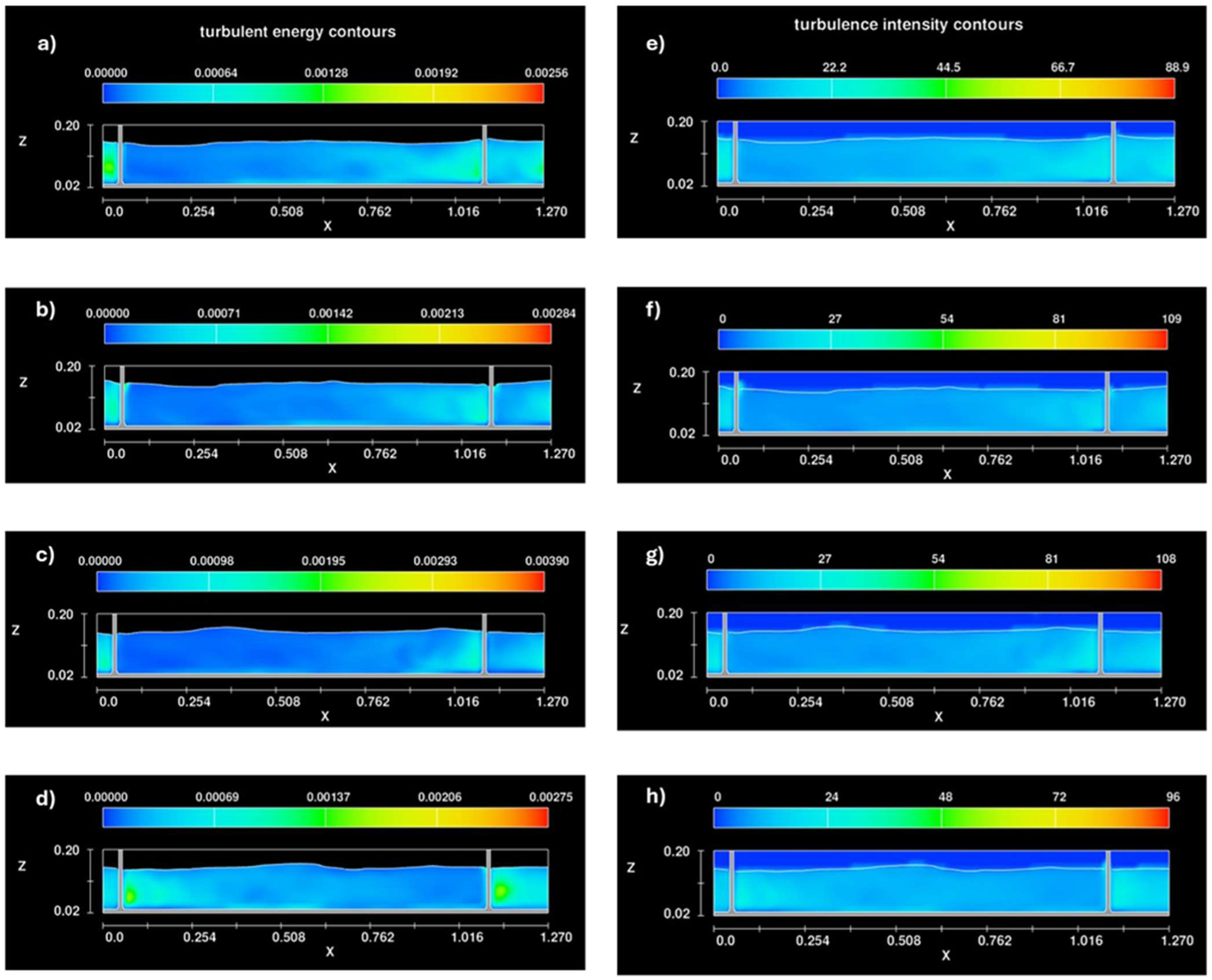
Wind generated turbulence energy (left), and intensity (right) predicted in the water column along the centre section of the flume in the simulations for (a, e) 7 knots, (b, f) 9 knots, (c, g) 11 knots and (d, h) 15 knots. Axes X and Z represent two-dimensions of the modelled flume chamber and results are measured in m^2^ s^-2^. Higher values of both turbulence energy and intensity indicate higher mixing and shear-induced stress on the gametes and embryos.

## 4. Discussion

As climate change alters global temperatures, corresponding wind regimes are predicted to change, often intensifying and promoting more frequent extreme conditions (Harley et al. 2006). Wind velocity predictions for the Great Barrier Reef (GBR) indicate that over the next century, changes will be variable across reefs, with stronger winds anticipated in the north and weaker winds in the south (McWhorter et al. 2022). Our study examined the impacts of potential changes in wind-driven surface mixing on fertilisation and embryogenesis of four species of corals found across the GBR, and latent larval survival effects for two of these species. Intact or non-defective embryos generally decreased with increased wind intensity and exposure time, owing to increases in damaged, deformed, and fragmented embryos. Damage was most common in the highest wind treatment (15 knots), and deformity occurred more frequently in the lower wind treatments (7, 9, 11 knots). In the 4 days following experimental exposure, intact *A. kenti* embryos had ∼3–fold and *A. spathulata* had ∼1.5–2-fold higher survival rates than damaged, deformed, or fragmented embryos. Thus, high wind spawning nights >8 knots may significantly reduce larval production, with the potential to disrupt system recovery, especially under degraded reef conditions and in windward reef locations.

Our results highlight different implications for survival across embryo condition categories that were consistent across species. Embryos in the most damaged states were the least likely to survive to the larval swimming stage. This has been observed in the literature for non-coral spawning invertebrates that were damaged due to laboratory turbulent mixing (Mead 1996). Fragmented embryos were able to survive, as corroborated by Heyward and Negri (2012), but survival rates were lower than normal embryos. Aside from normal embryos, deformed embryos had the highest likelihood of survival to the swimming stage of larval development. Randall and Szmant (2009) observed similar trends, where irregularly developing embryos were still able to survive to competency following minor heat stress. Okubo et al. (2017) found that experimentally fragmented embryos only survived when their animal pole was still intact. Animal poles have been demonstrated to be crucial for larval development in Cnidarians, especially during gastrulation and endoderm construction (Imai et al. 2000, Fritzenwanker et al. 2007). The animal pole also becomes the oral end of the larvae (Momose and Schmid 2006), which is required to maintain vital functions like feeding (Larsson et al. 2014). Therefore, the higher mortality rates of damaged and fragmented embryos in the present study may be a result of loss or damage of this integral animal hemisphere.

Species-specific variability in fertilisation has been documented in corals (Nozawa, Isomura and Fukami 2015, dela Cruz and Harrison 2020, Buccheri et al. 2023), likely owing to differences in gamete properties like egg size, sperm performance, recognition capabilities, and contact time between gametes (Levitan 2000, Levitan 2006, Morita et al. 2006, Nozawa, Isomura and Fukami 2015, Buccheri et al. 2023). Hydrodynamic mixing has also been observed to inhibit fertilisation success in other spawning invertebrates by diminishing sperm swimming speeds and sperm-egg binding capabilities, and these influences are variable across species (Mead 1996). Such trends were confirmed in the present study, with fertilisation and hydrodynamic sensitivity varying across species. *Acropora kenti* had the most rapid and highest fertilisation, supporting previous literature, which has recognised *A. kenti* for its fertility and efficiency compared to other species (Willis et al. 1997, Buccheri et al. 2023). *Acropora millepora*, *A. spathulata* and *P. daedalea* also had high fertilisation, but this was more sporadic and delayed than *A. kenti,* occurring ≥1 hour after mixing. Correspondingly, unfertilised eggs were more commonly observed with the latter three species, and within the first 90 min of trials. Unfertilised eggs are likely to be damaged as wind intensity and exposure time increases (Mead 1996) and deformed eggs can cause sperm chemotaxis to break down (Yoshida, Inara and Morisawa 1993), thus less efficient fertilisers will have less successful fertilisation overall when exposed to hydrodynamic stress for prolonged periods.

More nuanced variability was observed across all species in the other four condition categories, highlighting species-specific sensitivity to hydrodynamic stress. For example, *A. millepora* embryos were more likely to become deformed rather than damaged or fragmented. High levels of deformity have been reported for *A. millepora* in other studies, particularly when exposed to sedimentation (Humphrey et al. 2008). Fragmentation was most common in *P. daedalea*, and this was observed more frequently in the 11 and 15 knot treatments, while damage, deformity, and fragmentation were common in *A. spathulata* across all wind treatments after ∼90 mins. Variations in susceptibility are likely driven by differences in embryo size and structure across species (Mead 1996).

Coral reefs function within complex hydrodynamic systems that are driven by winds, waves, and their interactions with the water column and ocean bottom (Young and Hardy 1993, Huang et al. 2012, Lowe and Falter 2014, Harris et al. 2018, Harris et al. 2023). This study aimed to replicate windward shallow reef crest environments that experience spawning events during high wind conditions. These locations are prevalent along the GBR but are generally understudied due to suboptimal experimental conditions. In-tank hydrodynamic modelling from this study has supported previous findings that wave height, water velocity, and turbulence increase with increased wind speeds (Péquignet et al. 2011, Huang et al. 2012, Cheriton, Storlazzi and Rosenberger 2016). Regions with higher turbulent kinetic energy (*k*) have more energy available for breaking down flow structures and enhancing mixing and dispersion. Regions with higher turbulence intensity (*TI*) correspond to stronger mixing zones, while lower turbulence intensity indicates more stable or laminar-like regions.

Turbulence values from this study can be directly compared to *in* situ measurements, but there is little data available to do so currently on the GBR so future work should address this knowledge gap on reefs of interest. While our turbulence parameters *k* and *TI* are not routinely measured on coral reefs, turbulent energy dissipation (ε) can be approximated using 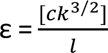 where *l* is the length scale, calculated as 3.8% of the water depth, and c is a constant equating to 0.9 (Rodi 1987, Burchard and Baumert 1995). Using our predicted turbulent kinetic energy (*k*) values, resulting mean turbulent energy dissipation (ε) in this study ranged from 1.6 × 10^-3^—3.7 × 10^-3^ W kg^-1^ from the 7- to 15-knot wind treatments, respectively, with peaks recorded >10^-2^ W kg^-1^ in the highest wind treatment (15 knots). These ε values are comparable to that observed in shallow windward coral reef lagoon on the GBR (10^-4^–10^-3^W kg^-1^) (Huang et al. 2012) but fall below the upper values reported for wave-exposed rocky shores (Gaylord 2008). In contrast, our ε values are at the upper ranges reported in studies reviewed by MacKenzie and Legget (1993) for wind-driven turbulence. This indicates that turbulence in our tanks is likely driven by both wind and wave forcing.

Beyond characterising the physical environment, our findings reveal a potential sensitivity of coral embryos to turbulent forces. We observed deformation during fertilisation at ε values markedly lower than those reported to cause similar deformation in urchin embryos in wave-exposed environments (Denny, Nelson and Mead 2002, Gaylord 2008). This difference suggests that coral embryos, which are known to fragment (Heyward and Negri 2012), might be more sensitive to turbulent fluid forces than other marine invertebrates like urchin embryos.

Overall, few studies have quantified turbulence on coral reefs (Davis, Pawlak and Monismith 2021, Norris et al. 2023), yet it is still well known that turbulence is driven by structural complexity and promotes mixing at local scales (Norris et al. 2023). In shallow reef systems, the residence times of water and planktonic particles are directly driven by local hydrodynamics (Monismith 2007, Reid et al. 2020). Thus, localised mixing will directly influence the flux of gametes during spawning events, and the quantity of competent larvae remaining following stress on spawning nights.

In deeper reef environments, outside of the scope of this study, intensified winds will have more severe impacts on fertilisation success and larval output (Mumby et al. 2024). Wind intensifies vertical mixing of gametes (Wolanski, Burrage and King 1989, Hamner, Colin and Hamner 2007), thus increases the likelihood of dilution before fertilisation can occur (Oliver and Babcock 1992, van Woesik 2010, Sakai et al. 2020). The present study conveys a more optimistic scenario of shallow reef environments where dilution does not occur, to better quantify post-fertilisation effects on embryogenesis and larval survival.

Our results highlight the capabilities of four common spawning corals to fertilise successfully in suboptimal wind-affected conditions, and the varying degrees of embryo sensitivity to damage, deformity, and fragmentation over time. Wind data are routinely available at the GBR scale, thus can be used to modify expectations for fertilisation and larval survival following exposure. Such results can inform models of coral reproduction, larval dispersal, and population dynamics (Graham, Baird and Connolly 2008, Graham et al. 2013, Teo and Todd 2018, Boschetti et al. 2020, Bozec et al. 2022, Figueiredo et al. 2022). Robust demographic metrics can help identify more realistic key source reefs (Mumby, Mason and Hock 2021), and will promote more appropriate management and restoration outcomes that safeguard natural reef recovery, given local abiotic conditions.

## Acknowledgements

The authors would like to acknowledge the Traditional Owners of the Great Barrier Reef, particularly the Byelle, Gooreng Gooreng, Gurang and Taribelang Bunda First Nations people and the Wulgurukaba and Bindal First Nations people for permission to collect corals from their Sea Country and bring them to laboratory facilities for experimentation. We would like to thank Jacqui Lardner, Rebecca Reeves, Mark Tonks, Frank Coman, Julian Uribe-Palomino, Robert Mason, Melanie Orr, Anthea Donovan, and Rosanna Griffith-Mumby for field support, Andrew Negri and Andrew Heyward for advice regarding the experiment, and the staff at Heron Island Research Station and the National Sea Simulator facility for assistance.

## CRediT authorship contribution statement

**E. B.** Conceptualisation, Methodology, Investigation, Data curation, Formal analysis, Visualisation, Writing - original draft, Writing - review and editing**. A. W.** Methodology, Software, Investigation, Data curation, Formal analysis, Visualisation, Writing - original draft, Writing - review and editing**. G. F. R.** Conceptualisation, Methodology, Formal analysis, Writing - review and editing, Supervision**. P. J. M.** Conceptualisation, Methodology, Writing - review and editing, Supervision, Funding acquisition**. C. D.** Conceptualisation, Methodology, Writing - review and editing, Supervision, Funding acquisition.

## Data statement

The datasets and code from the experimental component of this study are deposited in the CSIRO Data Access Portal http://hdl.handle.net/102.100.100/489500?index=1 and will be made fully available following publication.

