## Supplementary materials for "Impacts of wind-driven hydrodynamics on the early stages of coral development"

### Methods

#### *Hydrodynamic modelling*

FLOW-3D HYDRO is a computational fluid dynamics model that employs numerical techniques for solving the governing equations to obtain three-dimensional solutions for fluid flow problems. The model solves the following mass continuity (Eq. 1) and momentum equations (Eq. 2-4) in cartesian coordinates:

$$\frac{\partial}{\partial x}(uA_x) + R\frac{\partial}{\partial y}\left(vA_y\right) + \frac{\partial}{\partial z}(wA_z) + \frac{uA_x}{x} = \frac{R_{SOR}}{\rho} \quad (1)$$

$$\frac{\partial u}{\partial t} + \frac{1}{V_F}\{uA_x\frac{\partial u}{\partial x} + vA_yR\frac{\partial u}{\partial y} + wA_z\frac{\partial u}{\partial z}\} - \frac{A_y v^2}{xV_F} = -\frac{1}{\rho}\frac{\partial p}{\partial x} + G_x + f_x - b_x - \frac{R_{SOR}}{\rho V_F}(u - u_w - \delta u_s) \quad (2)$$

$$\frac{\partial v}{\partial t} + \frac{1}{V_F}(uA_x\frac{\partial v}{\partial x} + vA_yR\frac{\partial v}{\partial y} + wA_z\frac{\partial v}{\partial z}) + \frac{A_y uv}{xV_F} = -\frac{1}{\rho}R\frac{\partial p}{\partial y} + G_y + f_y - b_y - \frac{R_{SOR}}{\rho V_F}(v - v_w - \delta v_s) \quad (3)$$

$$\frac{\partial w}{\partial t} + \frac{1}{V_F}(\left\{uA_x\frac{\partial w}{\partial x} + vA_yR\frac{\partial w}{\partial y} + wA_z\frac{\partial w}{\partial z}\right\}) = -\frac{1}{\rho}\frac{\partial p}{\partial x} + G_z + f_z - b_z - \frac{R_{SOR}}{\rho V_F}(w - w_w - \delta w_s) \quad (4)$$

where the velocity components *u*, *v*, *w*, are in *x*, *y*, *z* directions respectively. $A_x$, $A_y$ and $A_z$ are the fractional areas open to flow in x, y, and z directions respectively and $V_F$ is the fractional volume open to flow. $R_{SOR}$ is a density source term (to model mass injection through porous medium, etc.), ρ is the fluid density, p is the pressure, ($G_x$, $G_y$, $G_z$) are body accelerations, ($f_x$, fy, fz) are viscous accelerations, ($b_x$, $b_y$, $b_z$) are flow losses. The governing equations are discretised using finite-volume approximations onto a regular grid of rectangular cells. An explicit solver is used to compute the quantities from the approximated conservation laws in Eq. 1-4. In addition to modelling the hydrodynamics, the model also includes a 'Particles' module to track the motion of particles in the flow.

***Turbulence model evaluation***

Turbulence is the natural unstable motion of fluids when the viscous forces are insufficient to stabilize fluid particle motion. This instability of fluid motion increases with increasing Reynolds numbers and results in formation of eddies. The turbulent fluid velocities in all the three directions are described by the quantities $u'$, $v'$ and $w'$. To describe this erratic motion of the fluid particles and dissipation, turbulence models are implemented in FLOW3D-HYDRO along with the previously described equations of fluid motion. In FLOW3D-HYDRO, there are four turbulence models available when setting up a 3D simulation: the two-equation $k$-$\varepsilon$ (T1 hereafter), Renormalization Group (RNG) (T2 hereafter), $k$-$\omega$ (T3 hereafter), and Large Eddy Simulation (LES) (T4 hereafter).

In the T1 model, the turbulent kinetic energy ($k$) and turbulence dissipation rate ($\varepsilon$) are calculated to update the eddy viscosity in the equation of fluid motion. The T2 turbulence model is similar to the former, except the equation constants found empirically in the $k$-$\varepsilon$ model are derived explicitly. The specific turbulent dissipation rate ($\omega$) is calculated instead of $\varepsilon$ in the T3 model. The T4 model resolves large-scale turbulent structures directly and approximates small scale features through an eddy viscosity quantity, thus, the T4 model requires high computational resources to capture turbulent structures with fine mesh resolutions.

***Model configuration and wind shear stress***

The simulation of wind-shear induced currents in a recirculating flume is a difficult task to replicate in CFD models. Common numerical modelling packages like FLOW3D-HYDRO, ANSYS or OpenFOAM have limitations that impose uniform wind on both the compartments of recirculating flumes (in the same direction, which creates opposing currents in the two compartments of the flume), thus circulation cannot be observed.

Initially, the configuration of the physical model flume as illustrated in Fig. 2 was replicated in FLOW3D-HYDRO (Fig. S2). The flume was set up as a rectangular tank of length 1.27m and width 0.4m, with a divider along the centre line to create two compartments. At the right and left end of the flume, curved dividers were created to direct the water current around the divider. The wind velocity was imposed on the water surface using 'Two-phase' flow physics module in combination with the 'Air entrainment' module. The wind imposed on top of the water surface did not create any current in the water column due to lack of wind shear stress on the water surface (within the 'Two-Phase' flow module). Therefore, the 'Wind' physics module was later chosen in which the wind shear stress is forced on the water surface using the formulation in Eq. 5, where $\rho_{air}$ is the density of air, $W$ is the wind speed above water surface, $C_D$ is the wind shear coefficient with a default value of 0.003. The unidirectional wind shear stress is applied uniformly over the water surface.

$$\tau_{s=\rho_a C_D |W| W}$$

(5)

The recirculation with two compartments in the physical model could not be reproduced due to the unidirectional model limitations of FLOW3D-HYDRO (and other CFD models). To address this, we used a novel approach to simulate the wind shear stress on the water surface and the resulting circulation in the flume. A rectangular domain of length 1.27 m, width 0.2 m and depth 0.2 m (representing only one compartment) was configured in the model assuming symmetry along the centre partition of the recirculating flume. The initial water depth of 0.12 m was specified similar to the laboratory experiments. The circulation was introduced in the model by imposing 'periodic' boundary conditions (included in FLOW3D-HYDRO) on either end of the domain such that the wind is forced from the left-hand boundary. The use of this 'periodic' boundary condition eliminates the requirement of two compartments for circulation of the current. The fluid and the particles leaving the right-hand boundary are reintroduced on the left-hand boundary, ensuring continuous circulation of the particles and the fluid in the flume. The lateral sides of the flumes were treated as walls so that no flow could occur through the lateral boundaries. The bottom of the flume was a rectangular solid of thickness 3 cm. The initial fluid in the domain was specified at an elevation of 0.15 m. All scenarios simulated with different wind speeds over the water surface are listed in Table S1.

In addition to these simulations, simulations were also performed with two cylindrical objects in the domain one at either end, to induce turbulence and lateral mixing of the flow, to replicate the turbulence and mixing observed in the physical model due to the dividers and curved edges on both ends (Fig. S3).

***Particle physics***

In the current simulations, the coral eggs and sperm were represented by 'mass particles' which simulates motion of suspended solids in the domain. The mass particles are defined by a set of properties such as diameter and density. In the current simulations, the following properties were prescribed for the solid particles as two different species.

FLOW-3D HYDRO models the motion of particles at a sub-grid scale using a Lagrangian physics model to track the motion and interaction of spherical particles that could be either solid mass particles or spherical droplets of fluid particles, with walls, fluid, or void regions. The dynamics of the mass particles is governed by the following equation of motion, assuming each particle is a sphere of diameter *d*.

$$\frac{du_p}{dt} = -\frac{1}{\rho_p}\nabla P + g + \beta\left(u - u_p\right)\left|u - u_p\right|.\frac{\rho}{\rho_p} + \frac{m_{added}}{m_p}\left(\frac{du}{dt} - \frac{du_p}{dt}\right)$$

(Eq. 6)

The equations of motion for the fluid and the particles are solved simultaneously to couple the momentum together. When the diffusion was enabled, an increment in position is provided to the particles which is evaluated by a Monte Carlo technique and added to its mean trajectory. The resistance on the particles is calculated using a drag force formulation such as $C_D = \left(\frac{24}{Re} + \frac{6}{1+\sqrt{Re}} + 0.4\right)$ where $C_D$ is the drag coefficient (=1 for sphere) and *Re* is the flow Reynolds number as discussed previously.

The motion of the mass particles was affected by the flow but the impact of mass particles on the flow itself is turned off by default. It is also important to note that diffusion of particles was turned off in the current simulations.

***Model configuration and wind shear stress***

The coral egg and sperm particles were introduced in the flow as spherical solids of diameter 400 µm and 40 µm, respectively, to simulate biologically relevant and logistically feasible *Acropora spp.* gametes (Babcock et al. 2003, Madin et al. 2016). The density of egg particles was defined as 1022.987 kg/m$^3$ (G.F. Ricardo personal comm.) and sperm were assumed to have neutral density. The number of eggs introduced in the flow was 3,000, whereas 50,000 sperm were introduced, to achieve roughly $10^4$ sperm $mL^{-1}$ concentration which has proven to be adequate for fertilisation across species (Nozawa, Isomura and Fukami 2015, dela Cruz and Harrison 2020, Buccheri et al. 2023). In the physical model, 50,000 sperm $mL^{-1}$ of the fluid was used. Only 50,000 sperm particles were used in the present simulations due to the computational constraints in simulating this quantity of particles. In the initial distribution of the particles a cluster of eggs (1,500) and sperm (25,000) were initially placed on two ends of the flume (Fig. S4). The particles were initially placed at uniform spacing throughout the water column with sperm and eggs separately (this distribution is referred to as D1 hereafter). To test the influence of the initial distribution of the particles in the water column, a second set of simulations were also performed with the particles initially positioned only in the top 2 cm of the water column, and with sperm and eggs together, to simulate gametes from individual colonies more accurately as illustrated in Fig. S5 (this distribution is represented as D2 hereafter). To improve lateral mixing in the numerical flume, simulations with cylindrical obstructions (as illustrated in Fig. S2) were also performed. The resulting dispersion of the particles in the flume was investigated and compared with that of earlier configuration of the numerical flume. All the simulations were run for a duration of 5 min after which no change in the particle distribution across the flume was observed.

### Results

#### *Influence of turbulence model*

The wind generated wave surface elevations in the tank were dependent on the turbulence model selected (Figure S6). The highest surface elevations were observed for turbulence model T4 while the surface elevations increased for models T1, T2 and T3, respectively. The flow velocities were correspondingly higher for T4 compared to the other models, as illustrated in Fig. S7. The particle dispersion in the fluid in turn was dependent on the choice of turbulence model. For models T1 to T3, the particles begin displacement with the commencement of wind shear on the water surface. After a few seconds, the particles were dispersed throughout the water column as illustrated in Fig. S7. There was a gap between the centre line of the flume and the particles in the T1 model. However, this gap reduced for T2, T3 and T4 models, with no distinct gap pattern observed for the T4 model.

***Influence of initial particle distribution***

The initial particle distribution had minimal impact on the final dispersion of the particles in the fluid. The flow velocities and the distribution of the particles at time t=500s are illustrated in Figure S8. The particle distribution across the flume was identical to that observed in Figure S7. Hence the initial distribution had minimal impact on the final distribution.

Overall, it was observed that the circulating flow characteristics in the simulations were highly dependent on the turbulence model, cylindrical obstructions, and the wind speed. However, all the configurations resulted in distribution of the eggs and sperm throughout the water column within a few seconds of the simulation time. While some turbulence models may take longer for mixing, others achieve it quickly. This highlights that more readily available turbulence model in CFD such as k-ε (T1) is sufficient to model this phenomenon at low computation cost compared to LES model (T4).

***Influence of cylindrical obstructions***

Flow separation and wrapping around the cylindrical objects were also observed. The water surface elevation with the introduction of the cylindrical objects is illustrated in Fig. S3.

### Tables

Table S1. List of numerical simulations performed in this study and descriptions of each of their input parameters. Wind speed was held constant in simulations 1-12 and was varied in 13-15. Two egg and sperm distribution options were examined, and four turbulence models were tested. Cylindrical particles were implemented to promote fertilisation in the latter half of the trials.

| Test number | Wind speed (m/s) | Egg distribution | Sperm distribution | Turbulence model | Cylindrical objects provided |
|---|---|---|---|---|---|
| 1 | 7.8 | D1 | D1 | T1 | No |
| 2 | 7.8 | D1 | D1 | T2 | No |
| 3 | 7.8 | D1 | D1 | T3 | No |
| 4 | 7.8 | D1 | D1 | T4 | No |
| 5 | 7.8 | D2 | D2 | T1 | No |
| 6 | 7.8 | D2 | D2 | T2 | No |
| 7 | 7.8 | D2 | D2 | T3 | No |
| 8 | 7.8 | D2 | D2 | T4 | No |
| 9 | 7.8 | D2 | D2 | T1 | Yes |
| 10 | 7.8 | D2 | D2 | T2 | Yes |
| 11 | 7.8 | D2 | D2 | T3 | Yes |
| 12 | 7.8 | D2 | D2 | T4 | Yes |

| | | | | | |
|---|---|---|---|---|---|
| 13 | 4.4 | D2 | D2 | T1 | Yes |
| 14 | 4.7 | D2 | D2 | T1 | Yes |
| 15 | 5.5 | D2 | D2 | T1 | Yes |

Table S2. Outputs from the alternative Dirichlet regression models examining the interactive influence of wind intensity and duration of exposure on the six main categories of embryo condition: fertilised, damaged, deformed, fragmented with cell divisions, fragmented without cell divisions, and unfertilised, for four species of spawning corals. Treatment 1 was used as an intercept in the models and the unfertilised egg classification was used as the reference category, thus both were omitted.

| **Species** | **Variable** | **Coefficient** | **Estimate** | **Std. Error** | **Z value** | **Pr(>\|z\|)** | **Corrected P** |
|---|---|---|---|---|---|---|---|
| | Fertilised | Treatment2 | 0.088 | 0.662 | 0.133 | 0.894 | 1.000 |
| | Fertilised | Treatment3 | -0.166 | 0.692 | -0.240 | 0.810 | 1.000 |
| | Fertilised | Treatment4 | -0.190 | 0.675 | -0.282 | 0.778 | 1.000 |
| | Fertilised | Time_since_mix | 0.004 | 0.004 | 1.005 | 0.315 | 1.000 |
| | Fertilised | Treatment2: Time_since_mix | <0.001 | 0.006 | 0.064 | 0.949 | 1.000 |
| ***A. kenti*** | Fertilised | Treatment3: Time_since_mix | <0.001 | 0.007 | 0.060 | 0.952 | 1.000 |
| | Fertilised | Treatment4: | 0.002 | 0.007 | 0.286 | 0.775 | 1.000 |

| | | | | | | | |
|---|---|---|---|---|---|---|---|
| | | Time_since_mix | | | | | |
| | Damaged | Treatment2 | 0.833 | 0.884 | 0.942 | 0.346 | 1.000 |
| | Damaged | Treatment3 | 0.269 | 0.911 | 0.295 | 0.768 | 1.000 |
| | Damaged | Treatment4 | 0.633 | 0.855 | 0.740 | 0.460 | 1.000 |
| | Damaged | Time_since_mix | 0.010 | 0.006 | 1.707 | 0.088 . | 1.000 |
| | Damaged | Treatment2: Time_since_mix | -0.003 | 0.008 | -0.306 | 0.759 | 1.000 |
| | Damaged | Treatment3: Time_since_mix | -0.001 | 0.009 | -0.135 | 0.893 | 1.000 |
| | Damaged | Treatment4: Time_since_mix | 0.003 | 0.008 | 0.369 | 0.712 | 1.000 |
| | Deformed | Treatment2 | -0.635 | 0.827 | -0.767 | 0.443 | 1.000 |
| | Deformed | Treatment3 | -0.876 | 0.845 | -1.037 | 0.300 | 1.000 |
| | Deformed | Treatment4 | -1.512 | 0.859 | -1.760 | 0.078 . | 1.000 |
| | Deformed | Time_since_mix | 0.003 | 0.005 | 0.582 | 0.561 | 1.000 |
| | Deformed | Treatment2: Time_since_mix | 0.006 | 0.008 | 0.726 | 0.468 | 1.000 |
| | Deformed | Treatment3: Time_since_mix | 0.010 | 0.008 | 1.222 | 0.222 | 1.000 |
| ***A. kenti*** | Deformed | Treatment4: Time_since_mix | 0.009 | 0.008 | 1.140 | 0.255 | 1.000 |
| | Fragmented with divisions | Treatment2 | 0.810 | 0.866 | 0.935 | 0.350 | 1.000 |
| | Fragmented with divisions | Treatment3 | 0.101 | 0.880 | 0.115 | 0.908 | 1.000 |
| | Fragmented with divisions | Treatment4 | 0.269 | 0.865 | 0.311 | 0.756 | 1.000 |

| | | | | | | | |
|---|---|---|---|---|---|---|---|
| | Fragmented with divisions | Time_since_mix | 0.005 | 0.006 | 0.929 | 0.353 | 1.000 |
| | Fragmented with divisions | Treatment2: Time_since_mix | -0.002 | 0.008 | -0.206 | 0.837 | 1.000 |
| | Fragmented with divisions | Treatment3: Time_since_mix | 0.004 | 0.008 | 0.515 | 0.606 | 1.000 |
| | Fragmented with divisions | Treatment4: Time_since_mix | 0.004 | 0.008 | 0.462 | 0.644 | 1.000 |
| | Fragmented no divisions | Treatment2 | 0.342 | 0.861 | 0.397 | 0.691 | 1.000 |
| | Fragmented no divisions | Treatment3 | 0.055 | 0.859 | 0.064 | 0.949 | 1.000 |
| | Fragmented no divisions | Treatment4 | 0.201 | 0.852 | 0.236 | 0.814 | 1.000 |
| | Fragmented no divisions | Time_since_mix | 0.008 | 0.005 | 1.424 | 0.155 | 1.000 |
| | Fragmented no divisions | Treatment2: Time_since_mix | -0.001 | 0.008 | -0.089 | 0.929 | 1.000 |
| | Fragmented no divisions | Treatment3: Time_since_mix | -0.002 | 0.008 | -0.276 | 0.782 | 1.000 |
| | Fragmented no divisions | Treatment4: Time_since_mix | -0.003 | 0.008 | -0.371 | 0.710 | 1.000 |
| ***A. spathulata*** | Fertilised | Treatment2 | 0.474922 | 0.713647 | 0.665 | 0.5057 | 1.000 |
| | Fertilised | Treatment3 | -0.859036 | 0.653049 | -1.315 | 0.1884 | 1.000 |
| | Fertilised | Treatment4 | 1.114150 | 0.739169 | 1.507 | 0.1317 | 1.000 |
| | Fertilised | Time_since_mix | 0.006368 | 0.004810 | 1.324 | 0.1856 | 1.000 |
| | Fertilised | Treatment2: Time_since_mix | -0.007188 | 0.006702 | -1.072 | 0.2835 | 1.000 |
| | Fertilised | Treatment3: Time_since_mix | 0.003057 | 0.006422 | 0.476 | 0.6341 | 1.000 |

| | | | | | | | |
|---|---|---|---|---|---|---|---|
| | Fertilised | Treatment4: Time_since_mix | -0.011599 | 0.006829 | -1.698 | 0.0894 . | 1.000 |
| | Damaged | Treatment2 | 1.195925 | 0.857027 | 1.395 | 0.162885 | 1.000 |
| | Damaged | Treatment3 | 0.055525 | 0.777194 | 0.071 | 0.943045 | 1.000 |
| | Damaged | Treatment4 | 1.486079 | 0.872361 | 1.704 | 0.088472 . | 1.000 |
| | Damaged | Time_since_mix | 0.020102 | 0.005352 | 3.756 | 0.000173 *** | 0.007 *** |
| | Damaged | Treatment2: Time_since_mix | -0.014139 | 0.007635 | -1.852 | 0.064049 . | 1.000 |
| | Damaged | Treatment3: Time_since_mix | -0.001181 | 0.007127 | -0.166 | 0.868339 | 1.000 |
| ***A. spathulata*** | Damaged | Treatment4: Time_since_mix | -0.013922 | 0.007648 | -1.820 | 0.068690 . | 1.000 |
| | Deformed | Treatment2 | 7.101e-01 | 8.257e-01 | 0.860 | 0.389755 | 1.000 |
| | Deformed | Treatment3 | -5.244e-01 | 8.088e-01 | -0.648 | 0.516786 | 1.000 |
| | Deformed | Treatment4 | 2.140e+00 | 8.574e-01 | 2.496 | 0.012576 * | 0.415 |
| | Deformed | Time_since_mix | 1.938e-02 | 5.253e-03 | 3.689 | 0.000225 *** | 0.009 *** |
| | Deformed | Treatment2: Time_since_mix | -1.007e-02 | 7.340e-03 | -1.372 | 0.169957 | 1.000 |
| | Deformed | Treatment3: Time_since_mix | 8.179e-06 | 7.375e-03 | 0.001 | 0.999115 | 1.000 |
| | Deformed | Treatment4: Time_since_mix | -2.261e-02 | 7.664e-03 | -2.950 | 0.003183 ** | 0.118 |
| | Fragmented with divisions | Treatment2 | 0.896194 | 0.866818 | 1.034 | 0.30119 | 1.000 |
| | Fragmented with divisions | Treatment3 | 0.244245 | 0.816952 | 0.299 | 0.76496 | 1.000 |
| | Fragmented with | Treatment4 | 1.130418 | 0.952087 | 1.187 | 0.23511 | 1.000 |

| | | | | | | | |
|---|---|---|---|---|---|---|---|
| | divisions | | | | | | |
| | Fragmented with divisions | Time_since_mix | 0.012434 | 0.005638 | 2.205 | 0.02743 * | 0.850 |
| | Fragmented with divisions | Treatment2: Time_since_mix | -0.008115 | 0.007814 | -1.039 | 0.29900 | 1.000 |
| | Fragmented with divisions | Treatment3: Time_since_mix | -0.003347 | 0.007643 | -0.438 | 0.66145 | 1.000 |
| | Fragmented with divisions | Treatment4: Time_since_mix | -0.012360 | 0.008551 | -1.445 | 0.14835 | 1.000 |
| ***A. spathulata*** | Fragmented no divisions | Treatment2 | 1.457590 | 0.834930 | 1.746 | 0.0809 . | 1.000 |
| | Fragmented no divisions | Treatment3 | -0.276086 | 0.781585 | -0.353 | 0.7239 | 1.000 |
| | Fragmented no divisions | Treatment4 | 0.201194 | 0.881315 | 0.228 | 0.8194 | 1.000 |
| | Fragmented no divisions | Time_since_mix | 0.012414 | 0.005519 | 2.249 | 0.0245 * | 0.784 |
| | Fragmented no divisions | Treatment2: Time_since_mix | -0.012067 | 0.007600 | -1.588 | 0.1124 | 1.000 |
| | Fragmented no divisions | Treatment3: Time_since_mix | 0.001975 | 0.007297 | 0.271 | 0.7867 | 1.000 |
| | Fragmented no divisions | Treatment4: Time_since_mix | -0.003605 | 0.007793 | -0.463 | 0.6437 | 1.000 |
| | Fertilised | Treatment2 | 0.086 | 0.653 | 0.131 | 0.895 | 1.000 |
| | Fertilised | Treatment3 | -1.190 | 0.622 | -1.912 | 0.056 . | 1.000 |
| | Fertilised | Treatment4 | -0.472 | 0.642 | -0.736 | 0.462 | 1.000 |
| ***A. millepora*** | Fertilised | Time_since_mix | 0.013 | 0.005 | 2.623 | 0.009 ** | 0.305 |
| | Fertilised | Treatment2: Time_since_mix | -0.001 | 0.007 | -0.167 | 0.867 | 1.000 |
| | Fertilised | Treatment3: Time_since_m | 0.010 | 0.007 | 1.501 | 0.133 | 1.000 |

| | | | | | | | |
|---|---|---|---|---|---|---|---|
| | | ix | | | | | |
| | Fertilised | Treatment4: Time_since_mix | 0.001 | 0.007 | 0.151 | 0.880 | 1.000 |
| | Damaged | Treatment2 | 0.056 | 0.796 | 0.071 | 0.944 | 1.000 |
| | Damaged | Treatment3 | -0.944 | 0.803 | -1.175 | 0.240 | 1.000 |
| | Damaged | Treatment4 | -0.351 | 0.818 | -0.430 | 0.668 | 1.000 |
| | Damaged | Time_since_mix | 0.015 | 0.006 | 2.484 | 0.013 * | 0.429 |
| | Damaged | Treatment2: Time_since_mix | <0.001 | 0.008 | 0.016 | 0.987 | 1.000 |
| | Damaged | Treatment3: Time_since_mix | 0.005 | 0.008 | 0.600 | 0.548 | 1.000 |
| | Damaged | Treatment4: Time_since_mix | -0.003 | 0.008 | -0.338 | 0.736 | 1.000 |
| | Deformed | Treatment2 | -0.201 | 0.836 | -0.241 | 0.810 | 1.000 |
| | Deformed | Treatment3 | -1.073 | 0.815 | -1.316 | 0.188 | 1.000 |
| | Deformed | Treatment4 | -0.263 | 0.816 | -0.322 | 0.747 | 1.000 |
| | Deformed | Time_since_mix | 0.017 | 0.006 | 2.704 | 0.007 | 0.254 |
| | Deformed | Treatment2: Time_since_mix | 0.004 | 0.008 | 0.488 | 0.625 | 1.000 |
| | Deformed | Treatment3: Time_since_mix | 0.011 | 0.008 | 1.310 | 0.190 | 1.000 |
| | Deformed | Treatment4: Time_since_mix | 0.002 | 0.008 | 0.194 | 0.846 | 1.000 |
| ***A. millepora*** | Fragmented with divisions | Treatment2 | 0.291 | 0.845 | 0.344 | 0.731 | 1.000 |
| | Fragmented with divisions | Treatment3 | -0.822 | 0.834 | -0.987 | 0.324 | 1.000 |

| | | | | | | | |
|---|---|---|---|---|---|---|---|
| | Fragmented with divisions | Treatment4 | 0.066 | 0.822 | 0.081 | 0.936 | 1.000 |
| | Fragmented with divisions | Time_since_mix | 0.017 | 0.006 | 2.673 | 0.008 ** | 0.271 |
| | Fragmented with divisions | Treatment2: Time_since_mix | -0.005 | 0.008 | -0.615 | 0.538 | 1.000 |
| | Fragmented with divisions | Treatment3: Time_since_mix | 0.006 | 0.008 | 0.759 | 0.448 | 1.000 |
| | Fragmented with divisions | Treatment4: Time_since_mix | -0.001 | 0.008 | -0.182 | 0.856 | 1.000 |
| | Fragmented no divisions | Treatment2 | -0.584 | 0.851 | -0.687 | 0.492 | 1.000 |
| | Fragmented no divisions | Treatment3 | -1.405 | 0.816 | -1.722 | 0.085 . | 1.000 |
| | Fragmented no divisions | Treatment4 | -0.553 | 0.807 | -0.685 | 0.493 | 1.000 |
| | Fragmented no divisions | Time_since_mix | 0.008 | 0.006 | 1.339 | 0.1806 | 1.000 |
| | Fragmented no divisions | Treatment2: Time_since_mix | 0.003 | 0.008 | 0.397 | 0.691 | 1.000 |
| | Fragmented no divisions | Treatment3: Time_since_mix | 0.012 | 0.008 | 1.428 | 0.153 | 1.000 |
| | Fragmented no divisions | Treatment4: Time_since_mix | 0.005 | 0.008 | 0.644 | 0.519 | 1.000 |
| ***P. daedalea*** | Fertilised | Treatment2 | -0.494 | 0.599 | -0.824 | 0.410 | 1.000 |
| | Fertilised | Treatment3 | -0.557 | 0.568 | -0.981 | 0.327 | 1.000 |
| | Fertilised | Treatment4 | 0.009 | 0.619 | 0.014 | 0.989 | 1.000 |
| | Fertilised | Time_since_mix | 0.015 | 0.005 | 2.958 | 0.003 ** | 0.105 |
| | Fertilised | Treatment2: Time_since_mix | 0.004 | 0.007 | 0.553 | 0.580 | 1.000 |

| | | | | | | | |
|---|---|---|---|---|---|---|---|
| | Fertilised | Treatment3: Time_since_mix | -0.002 | 0.006 | -0.394 | 0.693 | 1.000 |
| | Fertilised | Treatment4: Time_since_mix | -0.003 | 0.007 | -0.508 | 0.611 | 1.000 |
| | Damaged | Treatment2 | -0.407 | 0.805 | -0.506 | 0.613 | 1.000 |
| | Damaged | Treatment3 | -0.279 | 0.774 | -0.361 | 0.718 | 1.000 |
| | Damaged | Treatment4 | -0.046 | 0.810 | -0.057 | 0.955 | 1.000 |
| | Damaged | Time_since_mix | 0.017 | 0.006 | 2.869 | 0.004 ** | 0.136 |
| | Damaged | Treatment2: Time_since_mix | 0.001 | 0.008 | 0.077 | 0.938 | 1.000 |
| | Damaged | Treatment3: Time_since_mix | -0.005 | 0.008 | -0.593 | 0.553 | 1.000 |
| | Damaged | Treatment4: Time_since_mix | -0.002 | 0.008 | -0.184 | 0.854 | 1.000 |
| | Deformed | Treatment2 | -0.486 | 0.758 | -0.642 | 0.521 | 1.000 |
| | Deformed | Treatment3 | 0.686 | 0.714 | 0.960 | 0.337 | 1.000 |
| | Deformed | Treatment4 | 1.611 | 0.726 | 2.220 | 0.026 * | 0.811 |
| | Deformed | Time_since_mix | 0.018 | 0.006 | 3.102 | 0.002 ** | 0.069 |
| | Deformed | Treatment2: Time_since_mix | -0.001 | 0.008 | -0.154 | 0.878 | 1.000 |
| | Deformed | Treatment3: Time_since_mix | -0.013 | 0.007 | -1.710 | 0.087 . | 1.000 |
| | Deformed | Treatment4: Time_since_mix | -0.017 | 0.008 | -2.224 | 0.026 * | 0.811 |
| | Fragmented with divisions | Treatment2 | -1.163 | 0.828 | -1.403 | 0.161 | 1.000 |
| | Fragmented with | Treatment3 | -1.134 | 0.820 | -1.383 | 0.167 | 1.000 |

| | | | | | | | |
|---|---|---|---|---|---|---|---|
| | divisions | | | | | | |
| ***P. daedalea*** | Fragmented with divisions | Treatment4 | -0.213 | 0.788 | -0.270 | 0.787 | 1.000 |
| | Fragmented with divisions | Time_since_mix | 0.019 | 0.006 | 3.075 | 0.002 ** | 0.074 |
| | Fragmented with divisions | Treatment2: Time_since_mix | 0.009 | 0.008 | 1.120 | 0.263 | 1.000 |
| | Fragmented with divisions | Treatment3: Time_since_mix | 0.005 | 0.008 | 0.642 | 0.521 | 1.000 |
| | Fragmented with divisions | Treatment4: Time_since_mix | 0.006 | 0.008 | 0.716 | 0.474 | 1.000 |
| | Fragmented no divisions | Treatment2 | -0.050 | 0.846 | -0.059 | 0.953 | 1.000 |
| | Fragmented no divisions | Treatment3 | 0.012 | 0.791 | 0.016 | 0.987 | 1.000 |
| | Fragmented no divisions | Treatment4 | 0.817 | 0.825 | 0.990 | 0.322 | 1.000 |
| | Fragmented no divisions | Time_since_mix | 0.023 | 0.006 | 3.639 | <0.001 *** | 0.010 ** |
| | Fragmented no divisions | Treatment2: Time_since_mix | -0.004 | 0.009 | -0.422 | 0.673 | 1.000 |
| | Fragmented no divisions | Treatment3: Time_since_mix | -0.002 | 0.008 | -0.212 | 0.832 | 1.000 |
| | Fragmented no divisions | Treatment4: Time_since_mix | -0.007 | 0.008 | -0.833 | 0.405 | 1.000 |

Table S3. Summary of the survfit model examining the influence of embryo condition and the number of days following exposure on the survival of Acropora tenuis and A. spathulata embryo/larvae.

| Species | Condition category | Time | Number at risk | N.event | Survival rate | Std.error | Lower 95% CI | Upper 95% CI |
|---|---|---|---|---|---|---|---|---|
| *A. kenti* | Damaged | 1 | 104 | 11 | 0.894 | 0.030 | 0.837 | 0.955 |
| | | 2 | 78 | 13 | 0.745 | 0.045 | 0.661 | 0.840 |
| | | 3 | 52 | 16 | 0.516 | 0.057 | 0.415 | 0.641 |
| | | 4 | 26 | 17 | 0.179 | 0.052 | 0.101 | 0.316 |
| | Deformed | 0.5 | 95 | 4 | 0.958 | 0.021 | 0.918 | 0.999 |
| | | 1 | 76 | 7 | 0.870 | 0.037 | 0.800 | 0.945 |
| | | 2 | 57 | 7 | 0.763 | 0.050 | 0.671 | 0.867 |
| | | 3 | 38 | 10 | 0.562 | 0.066 | 0.447 | 0.707 |
| | | 4 | 19 | 11 | 0.237 | 0.070 | 0.133 | 0.421 |
| | Fragmented | 0.5 | 145 | 1 | 0.993 | 0.007 | 0.980 | 1.000 |
| | | 1 | 116 | 14 | 0.873 | 0.031 | 0.815 | 0.935 |
| | | 2 | 87 | 14 | 0.733 | 0.043 | 0.653 | 0.822 |
| | | 3 | 58 | 17 | 0.518 | 0.053 | 0.423 | 0.634 |
| | | 4 | 29 | 18 | 0.196 | 0.051 | 0.118 | 0.326 |
| | Normal | 1 | 72 | 3 | 0.958 | 0.024 | 0.913 | 1.000 |
| | | 2 | 54 | 1 | 0.941 | 0.029 | 0.885 | 0.999 |
| | | 3 | 36 | 2 | 0.888 | 0.045 | 0.804 | 0.981 |
| | | 4 | 18 | 3 | 0.740 | 0.087 | 0.589 | 0.931 |
| | Damaged | 0 | 333 | 6 | 0.982 | 0.007 | 0.968 | 0.996 |
| | | 1 | 279 | 17 | 0.922 | 0.016 | 0.892 | 0.953 |
| | | 2 | 224 | 36 | 0.774 | 0.026 | 0.724 | 0.827 |
| | | 3 | 168 | 53 | 0.530 | 0.033 | 0.469 | 0.599 |
| | | 4 | 112 | 55 | 0.270 | 0.030 | 0.217 | 0.336 |
| | | 5 | 56 | 55 | 0.005 | 0.005 | 0.001 | 0.034 |
| | | 0 | 167 | 1 | 0.994 | 0.006 | 0.982 | 1.000 |

| | | | | | | | | |
|---|---|---|---|---|---|---|---|---|
| *A. spathulata* | Deformed | 1 | 140 | 5 | 0.959 | 0.017 | 0.927 | 0.992 |
| | | 2 | 112 | 14 | 0.839 | 0.033 | 0.776 | 0.907 |
| | | 3 | 84 | 21 | 0.629 | 0.047 | 0.544 | 0.728 |
| | | 4 | 56 | 23 | 0.371 | 0.050 | 0.285 | 0.482 |
| | | 5 | 28 | 25 | 0.040 | 0.022 | 0.013 | 0.119 |
| | Fragmented | 1 | 185 | 10 | 0.946 | 0.017 | 0.914 | 0.979 |
| | | 2 | 148 | 23 | 0.799 | 0.031 | 0.740 | 0.863 |
| | | 3 | 111 | 33 | 0.561 | 0.041 | 0.486 | 0.648 |
| | | 4 | 74 | 35 | 0.296 | 0.039 | 0.228 | 0.383 |
| | | 5 | 37 | 36 | 0.008 | 0.008 | 0.001 | 0.056 |
| | Normal | 2 | 88 | 5 | 0.943 | 0.025 | 0.896 | 0.993 |
| | | 3 | 66 | 13 | 0.757 | 0.050 | 0.665 | 0.863 |
| | | 4 | 44 | 16 | 0.482 | 0.064 | 0.372 | 0.624 |
| | | 5 | 22 | 16 | 0.131 | 0.049 | 0.063 | 0.273 |

Table S4. Summary of the cox proportional hazards model testing the differences in survival between embryos from each condition category: damaged, deformed, and fragmented, compared to normal embryos for A. tenuis and A. spathulata.

| **Species** | **Condition category** | **Coefficient** | **Exp(coef)** | **Se(coef)** | **Lower 95% CI** | **Upper 95% CI** | **Z value** | **Pr(>\|z\|)** |
|---|---|---|---|---|---|---|---|---|
| *A. kenti* | Damaged | 1.6507 | 5.2109 | 0.3591 | 2.578 | 10.533 | 4.597 | <0.001*** |
| | Deformed | 1.5532 | 4.7266 | 0.3700 | 2.289 | 9.762 | 4.197 | <0.001*** |
| | Fragmented | 1.6477 | 5.1951 | 0.3563 | 2.584 | 10.445 | 4.624 | <0.001*** |

| | | | | | | | | |
|---|---|---|---|---|---|---|---|---|
| *A. spathulata* | Damaged | 0.7399 | 2.0958 | 0.1575 | 1.539 | 2.854 | 4.698 | <0.001*** |
| | Deformed | 0.4409 | 1.5540 | 0.1771 | 1.098 | 2.199 | 2.490 | 0.01 * |
| | Fragmented | 0.6594 | 1.9337 | 0.1661 | 1.396 | 2.678 | 3.971 | <0.001** |

**Figures**

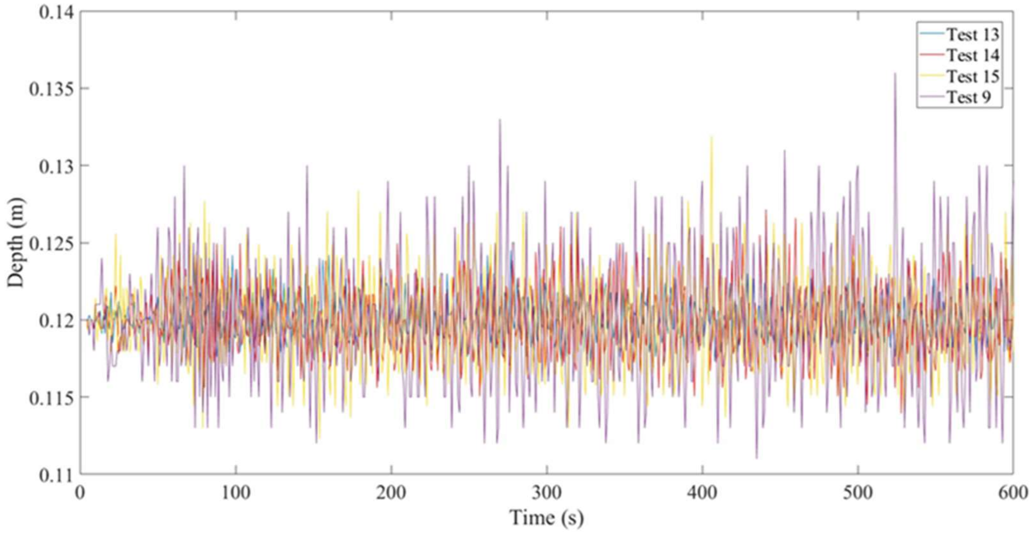


Fig. S1. Water surface elevations for different wind conditions in the T1 simulations, in the order of increasing wind speeds (a) Test 13 (7 knots), (b) Test 14 (9 knots), (c) Test 15 (11 knots) and (d) Test 9 (15 knots).

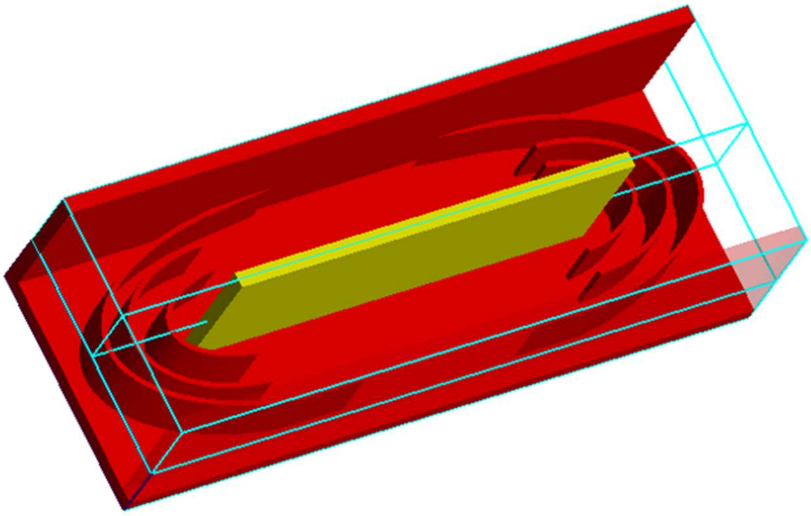

Figure S2. Configuration of the flume in the numerical model with two compartments separated by a divider wall and curved flow deflectors.

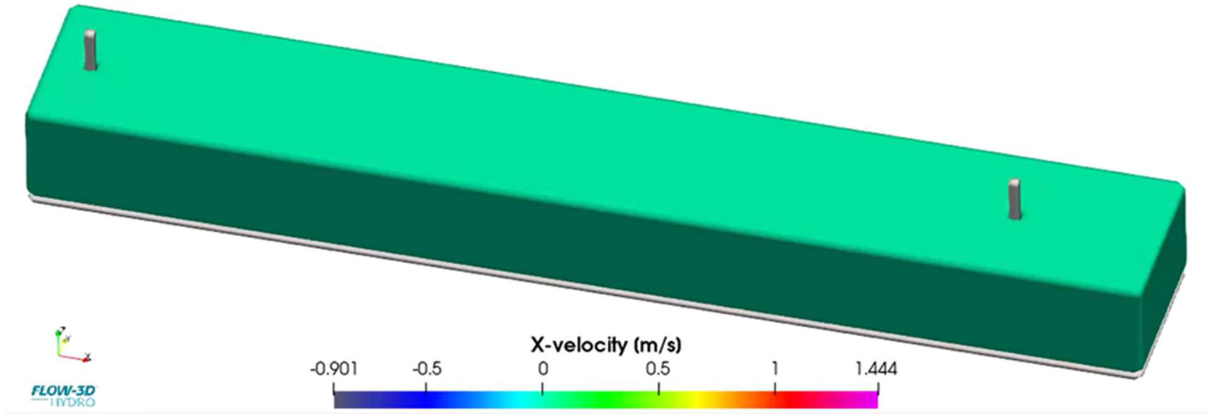


Figure S3. Configuration of the flume in the numerical model with a single compartment and two cylindrical objects one on either end. The initial fluid velocities are illustrated.

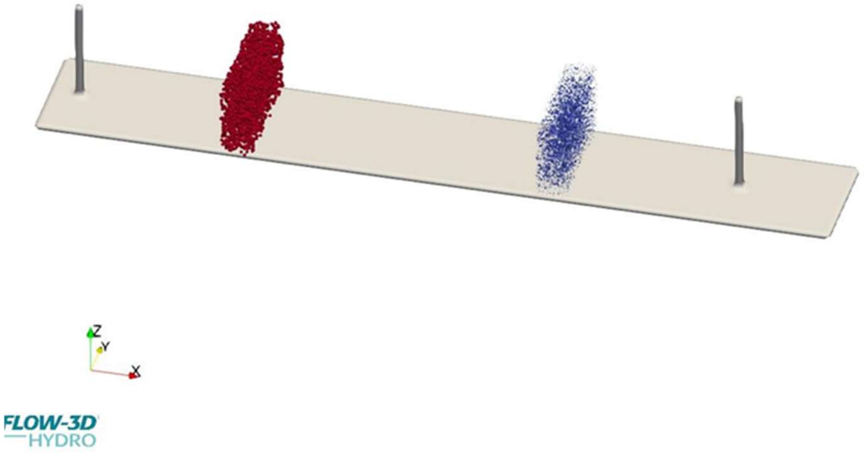

Figure S4. Illustration of initial particle distribution D1 in the flume (red – egg particles; blue – sperm particles). Particles were placed at each end of the flume tank separately to examine the feasibility of the model design, then was improved.

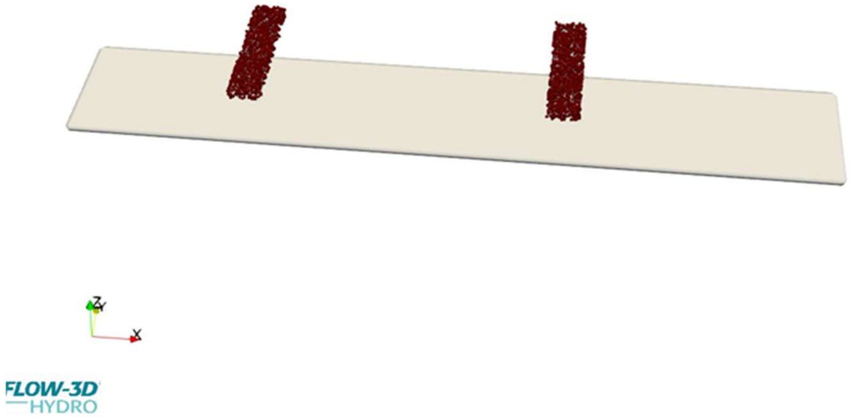

Figure S5. Illustration of initial particle distribution D2 in the flume, with egg and sperm particles introduced in both locations, to simulate eggs (red) and sperm (blue) from bundles more realistically.

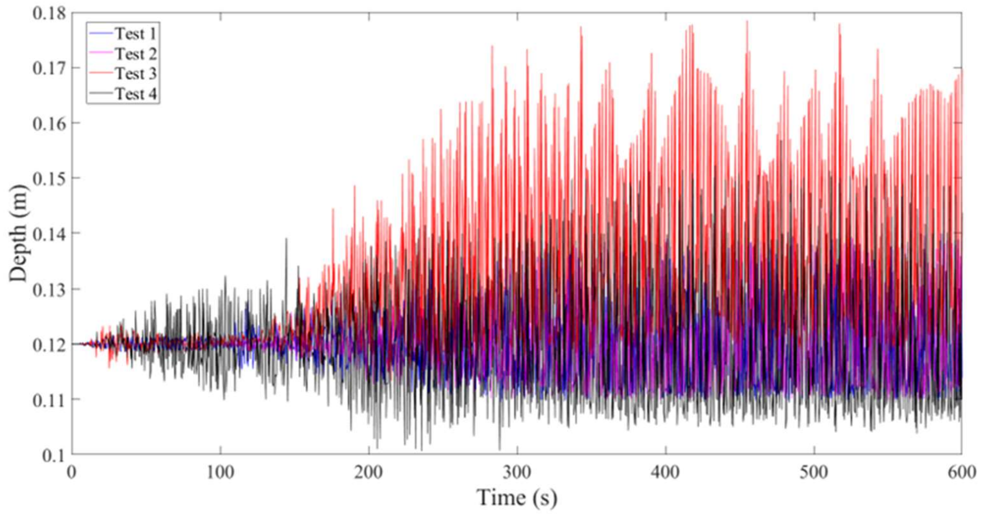


Figure S6. Water surface elevation for different turbulence models (a) Test 1, (b) Test 2, (c) Test 3, (d) Test 4.

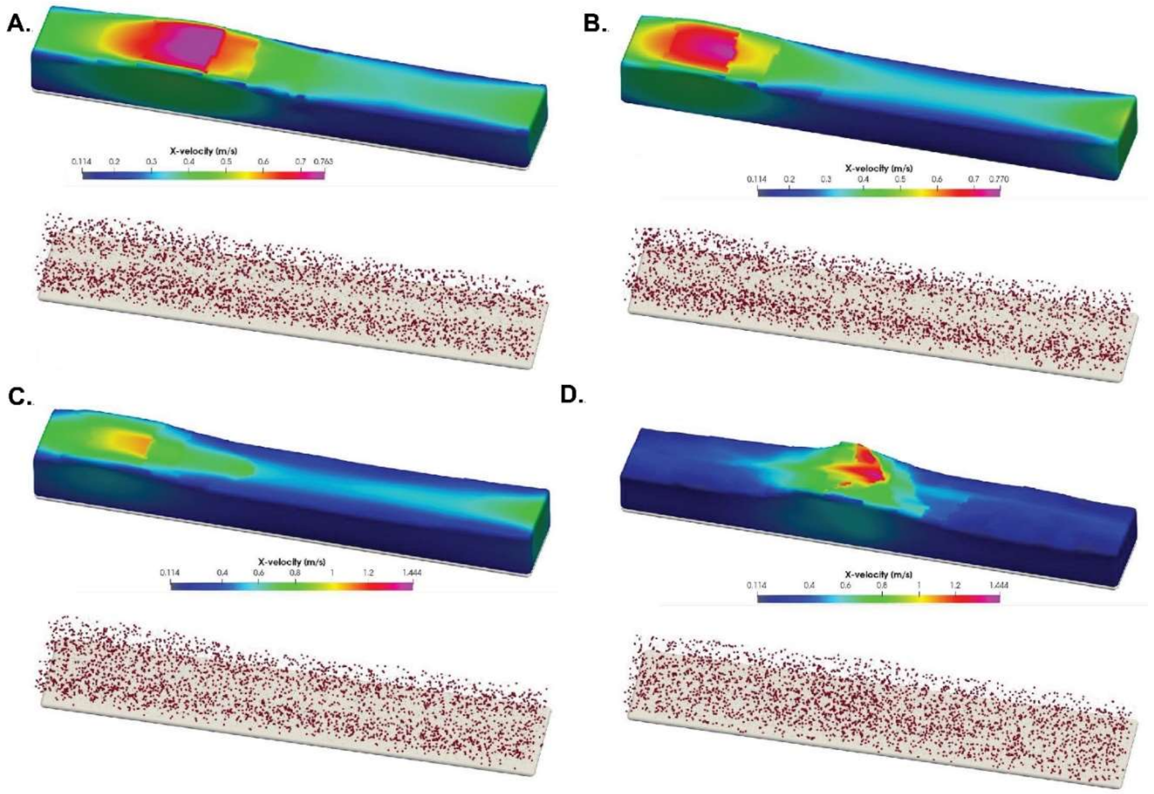


Figure S7. Fluid velocity contour map and particle distribution at time 500s across the different turbulence models (a) Test 1, (b) Test 2, (c) Test 3 and (d) Test 4.

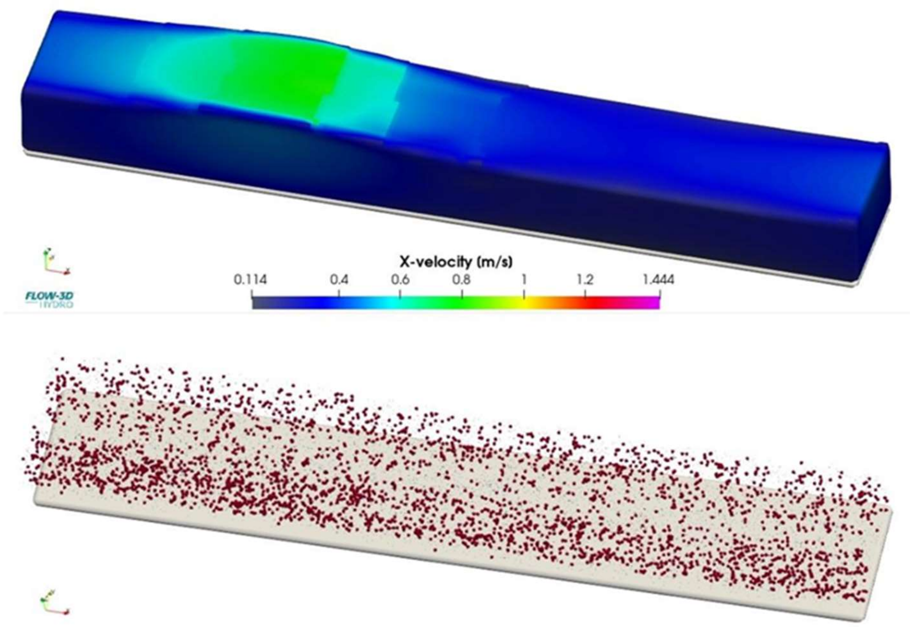


Figure S8. Fluid velocity contour map and particle distribution at time 500s in Test 5.
